# Spatial colocalization and combined survival benefit of natural killer and CD8 T cells despite profound MHC class I loss in non-small cell lung cancer

**DOI:** 10.1101/2024.02.20.581048

**Authors:** Remziye E. Wessel, Nardin Ageeb, Joseph M. Obeid, Ileana Mauldin, Kate A. Goundry, Gabriel F. Hanson, Mahdin Hossain, Chad Lehman, Ryan D. Gentzler, Nolan A. Wages, Craig L. Slingluff, Timothy N.J. Bullock, Sepideh Dolatshahi, Michael G. Brown

## Abstract

**Background:** MHC class I (MHC-I) loss is frequent in non-small cell lung cancer (NSCLC) rendering tumor cells resistant to T cell lysis. NK cells kill MHC-I-deficient tumor cells, and although previous work indicated their presence at NSCLC margins, they were functionally impaired. Within, we evaluated whether NK cell and CD8 T cell infiltration and activation vary with MHC-I expression.

**Methods:** We used single-stain immunohistochemistry (IHC) and Kaplan-Meier analysis to test the effect of NK cell and CD8 T cell infiltration on overall and disease-free survival. To delineate immune covariates of MHC-I-disparate lung cancers, we used multiplexed immunofluorescence (mIF) imaging followed by multivariate statistical modeling. To identify differences in infiltration and intercellular communication between IFNγ-activated and non-activated lymphocytes, we developed a computational pipeline to enumerate single cell neighborhoods from mIF images followed by multivariate discriminant analysis

**Results:** Spatial quantitation of tumor cell MHC-I expression revealed intra- and inter-tumoral heterogeneity, which was associated with the local lymphocyte landscape. IHC analysis revealed that high CD56^+^ cell numbers in patient tumors were positively associated with disease-free survival (DFS) (HR=0.58, *p*=0.064) and overall survival (OS) (HR=0.496, *p*=0.041). The OS association strengthened with high counts of both CD56^+^ and CD8^+^ cells (HR=0.199, *p*<1×10^-3^). mIF imaging and multivariate discriminant analysis revealed enrichment of both CD3^+^CD8^+^ T cells and CD3^−^CD56^+^ NK cells in MHC-I-bearing tumors (p<0.05). To infer associations of functional cell states and local cell-cell communication, we analyzed spatial single cell neighborhood profiles to delineate the cellular environments of IFNγ^+/-^ NK cells and T cells. We discovered that both IFNγ^+^ NK and CD8 T cells were more frequently associated with other IFNγ^+^ lymphocytes in comparison to IFNγ^−^ NK cells and CD8 T cells (p<1×10^-30^). Moreover, IFNγ^+^ lymphocytes were most often found clustered near MHC-I^+^ tumor cells.

**Conclusions:** Tumor-infiltrating NK cells and CD8 T cells jointly affected control of NSCLC tumor progression. Co-association of NK and CD8 T cells was most evident in MHC-I-bearing tumors, especially in the presence of IFNγ. Frequent co-localization of IFNγ^+^ NK cells with other IFNγ^+^ lymphocytes in near-neighbor analysis suggests NSCLC lymphocyte activation is coordinately regulated.

**What is already known on this topic:** MHC-I loss occurs frequently in NSCLC and corresponds with waning immunity in the tumor microenvironment (TME). NK cells recognize “missing-self” targets and could be leveraged to target NSCLC tumors with MHC-I loss. While NK cell presence at tumor margins has been documented in NSCLC, they were shown to lose function in this environment.

**What this study adds:** We developed spatial analysis pipelines leveraging the local heterogeneity of the TME at single cell resolution to test whether NK cells and T cells together contribute antitumoral immunity in NSCLC. We discovered that a high density of tumor-infiltrating NK cells corresponded with DFS, and this association was increased in patients with high coincident CD8 T cells, especially those in central tumor. Intriguingly, both cell types were found clustered together in MHC-I-bearing tumors, especially when both expressed IFNγ, suggesting coordinated lymphocyte activities may enhance immune control of NSCLC.

**How this study might affect research, practice, or policy:** This study provides a rationale for developing novel immunotherapies that simultaneously increase NK and T cell anti-tumoral immunity. Associations linking NK cells with patient survival and increased immune effector activity in NSCLC, even in MHC-I-deficient tumors, further highlights the need to devise and deploy NK cell activating strategies which may be highly efficacious in CD8 T cell refractory NSCLC.

**Graphical Abstract:** 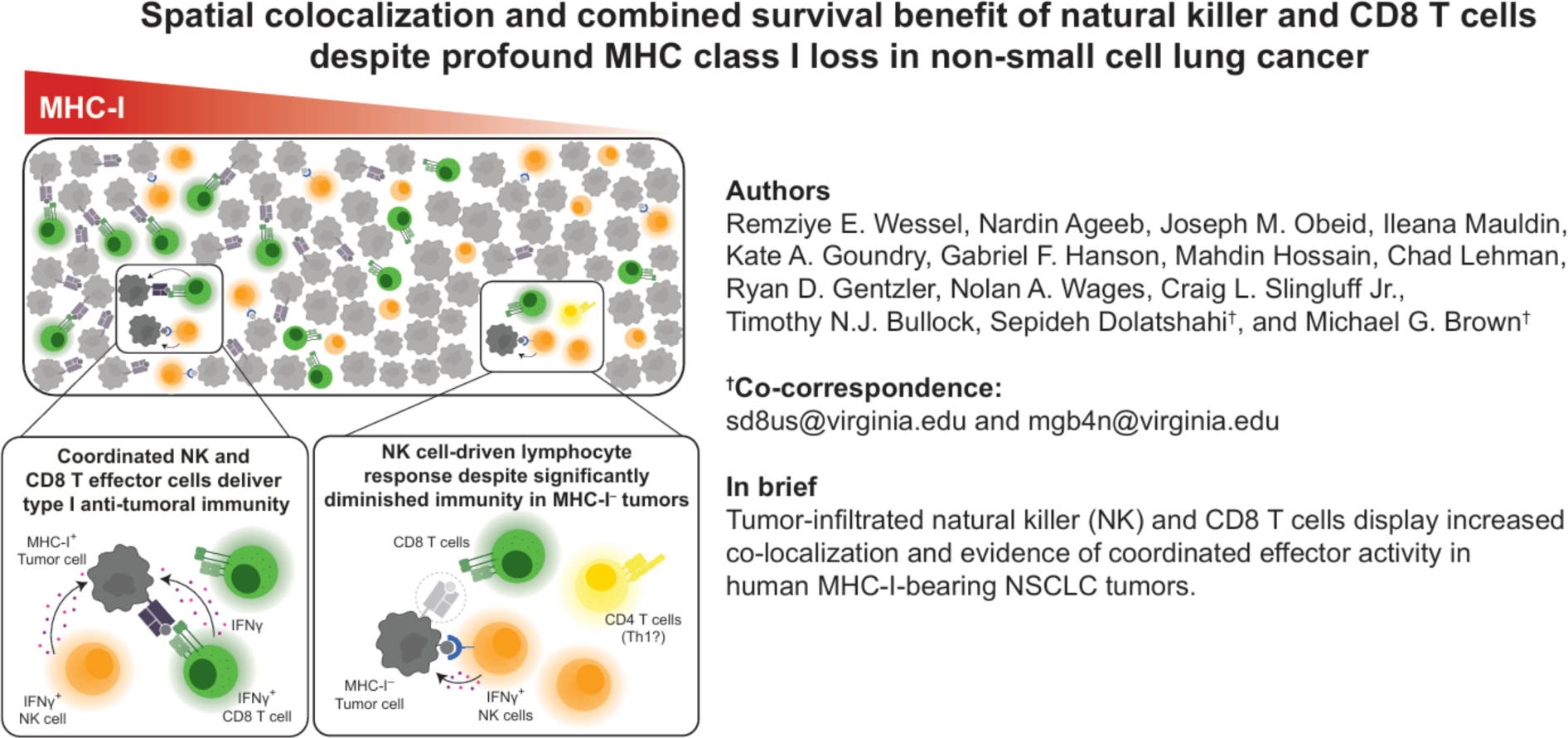

## BACKGROUND

Lung cancer is the leading cause of cancer-related death in the United States and globally, and NSCLC accounts for ∼85% of all lung cancers.^1^ ^2^ Adenocarcinoma (AdenoCA) and squamous cell carcinoma (SCC) together account for ∼90% of all NSCLC subtypes.^3^ ^4^ Both tumor types are infiltrated by diverse immune cells, including T cells and NK cells with cytokine and cytotoxic effector activity,^5^ highlighting the opportunity to harness these cytolytic cells to mediate efficacious control of lung cancer.

Tumor cell MHC-I loss to evade recognition by cytotoxic T cells is common in NSCLC, occurring in >30% of human lung cancer cells with >90% having defects of one or more MHC-I antigens or other molecules associated with antigen presentation.^6^ This constitutes a logical evolutionary step given the propensity of NSCLC to have a large repertoire of antigenic mutations that can be presented to T cells on MHC-I.^7^ Despite the frequency of MHC-I loss, its impact on lymphocyte infiltration and patient survival has not been resolved.^8–14^

The density of CD8 T cells infiltrating NSCLC tumors has been associated with a favorable prognosis.^15–17^ However, assessment of patient risk based on this determination alone is challenging due to factors such as the heterogeneity of intratumoral CD8 T cell infiltrates, the inherent difficulty in reliably sampling and assessing tumor-infiltrating lymphocytes, a plausible role for other immune cells, and because T cell exhaustion can contribute to poor immunological control of lung cancers.^18–20^ However, NK cells have been shown to provide protection against hematological malignancies or metastases from primary tumors, yet their antitumor activity has been less evident in solid tumors.^21^ ^22^ Given their ability to lyse ‘missing-self’ targets, NK cells represent a presently unharnessed, yet logical candidate to improve anti-tumor immunity in MHC-I-deficient, T cell-refractory lung tumors.^23^

Prior work has shown that NK cells are prevalent in β2-microglobulin (β2m-)deficient NSCLC tumors,^13^ but intratumoral NK cells from NSCLC patients were found to have reduced IFNγ production and cytolytic capability compared to circulating NK cells from the same patients. ^24^ Nonetheless, tumor-infiltrating NK cells activated by cytokines or stimulated via direct recognition of tumor cell expressed ligands of NK activating receptors have been shown to enhance tumor-specific T cells and control of tumor progression, possibly via cytokines to enhance recruitment or activation of classical dendritic cells (cDC).^25–30^ Understanding NK cell and CD8 T cell interactions supporting the anti-tumor immune response in relation to MHC-I expression in the NSCLC TME is a priority.

Hypothesizing that NK cells and CD8 T cells together drive anti-tumor immunity in NSCLC in relation to tumor cell MHC-I expression, we investigated whether tumor-infiltrating NK cells and CD8 T cells were associated with patient survival, and if tumor cell MHC-I expression corresponded with variations in lymphocyte presence, activation status, or spatial associations in the NSCLC TME. Our findings suggest that novel strategies designed to leverage both T cells and NK cells in combined immunotherapy may provide enhanced antitumor immunity and increased T cell activation, even in T cell refractory tumors.

## MATERIALS AND METHODS

### Clinical Patient Enrollment

Patients with non-small cell lung cancer (NSCLC) seen at the University of Virginia between 2011-2014 were enrolled in the clinical Cohort 1 (IRB-HSR#:18346). These participants had a resection at UVA in the years 2000 – 2014 and an available specimen in the UVA Biorepository and Tissue Research Facility. Patients were excluded only if the specimen was inadequate for testing (i.e., mostly fibrotic or necrotic tissue, or does not contain tumor cells). Clinical staging data is shown for all participants (**Table 1**), including those patients who underwent resection to non-evident disease.

**Table 1.** NSCLC patient characteristics.

|  |  | Cohort 1 (n=151) (Retrospective) |  |  | Cohort 2 (n=36) (Prospective) |  |  | Total (Cohort 1 + Cohort 2) |
| --- | --- | --- | --- | --- | --- | --- | --- | --- |
| Clinical feature * |  | AdenoCA (n=93) | SCC (n=58) | Total | AdenoCA (n=28) | SCC (n=8) | Total |  |
| Biologic sex | Female | 49 (53%) | 21 (36%) | 70 (46%) | 20 (71%) | 5 (63%) | 25 (69%) | 95 (51%) |
| Age in years, mean +/- (range) |  | 67 ± 9 (49-89) | 66 ± 9 (38-84) | 67 ± 9 (38-89) | 66 ± 11 (47-85) | 62 ± 12 (55-81) | 65 ± 11 (47-85) | 66 ± 9 (38-89) |
| AJCC v7 Stage at excision | Stage I | 57 (61%) | 36 (62%) | 93 (62%) | 7 (25%) | 0 (0%) | 7 (19%) | 100 (53%) |
|  | Stage II | 20 (22%) | 13 (22%) | 33 (22%) | 10 (36%) | 2 (25%) | 12 (33%) | 45 (24%) |
|  | Stage III | 13 (14%) | 8 (14%) | 21 (14%) | 11 (39%) | 5 (63%) | 16 (44%) | 37 (20%) |
|  | Stage IV | 3 (3%) | 1 (2%) | 4 (3%) | 0 (0%) | 1 (13%) | 1 (3%) | 5 (3%) |
| Largest Tumor Diameter* in cm, Mean +/- SD (range) |  | 2.7 ± 1.8 (0.6-12) | 3.7 ± 2.4 (0.9-10) | 3.1 ± 2.1 (0.6-12) | 3.29 ± 1.58 (1.3-6.3) | 3.03 ± 1.58 (1.2-5.5) | 3.23 ± 1.56 (1.2-6.3) | 3.12 ± 1.99 (0.6-12) |
| Prior chemotherapy |  | 5 (6%) | 1 (2%) | 6 (4%) | 6 (21%) | 3 (38%) | 9 (25%) | 15 (8%) |
| Treated with anti-PD(L)1 therapy |  | 0 (0%) | 0 (0%) | 0 (0%) | 4 (14%) | 0 (0%) | 4 (11%) | 4 (2%) |
| * number (%) unless otherwise specified |  |  |  |  |  |  |  |  |

### Immunohistochemistry

The AJCC 7^th^ edition was used for clinical and pathological staging. A pathologist outlined tumor areas and margins. Hematoxylin-eosin-stained sections from formalin-fixed, paraffin embedded (FFPE) tissues were used to identify regions of interest for analysis. Central tumor (CT) and peripheral tumor (PT) 1-mm cores, identified by a pathologist, were extracted from the FFPE tissues and used to construct the tissue microarray (TMA) as described.^31^ The TMA was incubated with primary antibodies to β2-microglobulin (mAb NAMB-1, kindly provided by Soldano Ferrone), CD3 (mAb Dako, clone 7.2.38), CD8 (clone C8/144B, Dako Carpinteria, CA), CD56 (mAb NCL-L-CD56-1B6, Leica), Granzyme B (ab4059, Abcam), and HLA class I heavy chains (HC) (HC10,^32^ kindly provided by Soldano Ferrone). Enzyme-linked secondary antibodies and 3,3’-diaminobenzidine (Dako) were used to develop the staining. Antibody-stained slides were counterstained with hematoxylin (Dako) and permanently mounted as described.^31^ Images were acquired using a Nikon Digital Sight DS-Fi1 camera (Nikon Corporation, Tokyo, Japan), an Eclipse 400 microscope (Nikon), or a NIS-Elements Viewer version 3.2 (Nikon). Control tissues included tonsil, lymph node, and placenta. Tissues incubated with PBS instead of primary antibody followed by staining with secondary antibody served as negative controls.

### TMA Immunotyping

Tumor cell antigen expression was assessed by a pathologist and scored by expression intensity (0=none, 1=mild, 2=strong). Scores from two cores per patient were summed to account for tumor heterogeneity. Automated CD8 counting was performed using a Leica SCN400 Scanner (Leica, Nussloch, Germany) and Digital Imaging Hub (Leica) and validated by manual counting by a pathologist. Immune cell counts were reported as (counts/mm^2^).

### Multiplex immunofluorescence (mIF) imaging and geospatial analysis

Five µm-thick sections from FFPE resected tumors were analyzed. Human lymph node samples were used as a positive control. mIF was performed according to the manufacturer’s protocol using the OPAL Multiplex Manual IHC kit and antigen retrieval (AR) buffers AR6 and AR9 (Akoya Biosciences, Marlborough, Massachusetts, USA) and DIVA Decloaker AR buffer (Biocare Medical, Pacheco, California, USA). Staining sequence, antibodies and AR buffers used are as follow: AR9, CD3 (1:100, Cell Marque MRQ-39) Opal520; AR9, CD8 (1:500, clone C8/144B Agilent Technologies, Santa Clara, California, USA) Opal540; AR6, CD56 (1:1000, Cell Marque MRQ-42) Opal570; DIVA, IFNγ (1:1000 NeoBiotechnologies 345 MSM4 P2 IFNG/466) Opal620; AR6, MHC-I (1:15,000 HLA class I heavy chains – HC10,^32^ Ferrone Lab) Opal650; AR6 Pan-cytokeratin (panCK) (1:200 MA513203, Invitrogen) Opal690, and AR6, spectral DAPI (Akoya Biosciences) as described.^33^

Stained slides were mounted using prolong diamond antifade (Life Technologies, Carlsbad, California, USA) and scanned using the PerkinElmer Vectra 3.0 system and Vectra software (Akoya Biosciences). Regions (3-mm^2^) were identified using Phenochart software and tissue images were acquired at 20x magnification with the Vectra 3.0 system. The images were spectrally unmixed using single stain positive control images using InForm software (Akoya Biosciences). A tissue classifier was trained on panCK^+^ tumor cells and panCK^−^ stroma followed by cell segmentation using HALO software (Indica Labs, Albuquerque, New Mexico, USA). Lymphocyte density (cells/mm^2^) was quantified for a given marker within stromal or tumor compartments of each region. To increase statistical power while avoiding size-related representational biases in resected tissues, a patient-agnostic, tissue regional analysis was developed to analyze scanned images of AdenoCA and SCC patient tumor biopsies as independent entities (**supplemental figure S1**).

### Spatial transcriptomics

The GeoMx Digital Spatial Profiler (DSP) (Nanostring) was used to generate whole transcriptome data for a subset of Cohort 1 patient tumors. FFPE tissue slices (5 μm) were sectioned onto positively charged slides. Regions of interest (ROIs) including at least 250 cells were selected based on staining with fluorophore-conjugated mAbs against panCK (AE-1/AE-3, Novus Biologicals), CD8 (C8/144B, Biolegend), and MHC-I HC (HC10, Ferrone Lab). Oligo-tagged profiling reagents were used to interrogate transcript expression within ROIs. Transcripts were read using an Illumina Sequencer and gene expression was quantified using the DSP interactive software based on the Whole Transcriptome Atlas.

### Survival analyses

Univariate and multivariate survival analysis was performed using R v.4.2.1. *Kaplan-Meier analysis*. To test the effect of NK and CD8 T cell abundances on OS (the time from resection to death by any cause) and DFS (the time from surgical resection to disease recurrence or death by any cause), patient tumors were stratified at an optimal cut point determined as described by Contal and O’Quigley ^34^ based on cell density (cells/mm^2^) averaged between CT and PT regions. The effect of MHC-I expression on OS and DFS was similarly determined by comparing patients with MHC-I expression above or below 60% averaged between CT and PT. Significance was determined by a log-rank test (*p*<0.05). Tumors with missing data for any of these markers were excluded from those respective analyses. *Cox proportional hazards modeling*. Proportional hazards assumptions were tested using Schoenfeld residual analysis for univariate and multivariate analysis (*p*<0.01). Multivariate Cox proportional hazards models assessed the relationship between survival endpoints (OS and DFS) and NK cell, CD8 T cell abundances, and MHC-I expression after adjusting for age and sex. Associations were considered significant for two-sided *p* values≤0.05.

### Cellular neighborhood analysis

*Custom cell-cell neighborhood scoring algorithm*. Spatial colocalizations were determined in Python using single cell 2-dimensional coordinates obtained from HALO. The Euclidian distance between each cell and every other cell on the slide was computed; nearest neighbors were defined as cells with a center-to-center Euclidean distance <30μm (**supplemental figure S2**). The nearest neighbors of each phenotype were enumerated to yield the neighborhood profile for each cell. *Intercellular K function analyses.* Intercellular geospatial colocalizations were further analyzed using the Spatstat package in R v.4.2.1.^35^ The K function was used to determine the number of nearest neighbors for a given cell type as a function of radius <200μm for each tumor region separately.

### Multivariate discriminant analyses

Orthogonalized Partial Least Squares Discriminant Analyses (OPLSDA) two-component models were generated in MATLAB (R2022a, MathWorks, Natick, MA) using scripts developed in-house.

All data input to OPLSDA were log-transformed, centered, and scaled. OPLSDA models were constructed on cell densities per region (cells/mm^2^) to discriminate between tumor regions with MHC-I^+^ tumor cell counts (panCK^+^MHC-I^+^) above or below the median for AdenoCA and SCC tumors separately. Cells localized to the tumor or stroma were normalized to their respective compartment areas. Second, OPLSDA models were constructed to discriminate the cellular neighborhood profiles between IFNγ^+/-^ NK cells (CD3^−^CD56^+^IFNγ^+/-^) and CD8 T cells (CD3^+^CD8^+^IFNγ^+/-^). Prediction accuracy was determined using a random 5-fold cross validation (CV) framework. Significance was determined empirically by comparing the model’s CV accuracy against 1000 randomly permuted models.^36^ ^37^ Univariate comparisons between model features were conducted using a non-parametric Wilcoxon rank sum test with a Benjamini-Hochberg false discovery rate correction controlled at *p*<0.05.

## RESULTS

### Patient characteristics

TMA specimens collected from patients seen at UVA between 2011-2014 were analyzed by single-stain immunohistochemistry (IHC) along with retrospective review of clinical records (IRB-HSR#:18346) (Cohort 1) (**table 1**). Whole tumor resections collected prospectively from patients seen at UVA between 2014-2018 were analyzed by mIF (IRB-HSR# 13310) (Cohort 2). Two patients in Cohort 2 were treated with Pembrolizumab, one was treated with Atezolizumab, and one was treated with Nivolumab following surgical resection. No patients in Cohort 1 received immunotherapy.

### MHC-I expression variation in human NSCLC is associated with variation in intra-tumoral lymphocytes

To examine MHC-I expression in NSCLC tumors and its relationship with occupation by CD56^+^ NK cells and CD8 T cells, we performed IHC on Cohort 1 (**table 1**) staining for anti-MHC-I HC, β2m, CD56, and CD8. We observed wide-ranging MHC-I HC expression levels in both AdenoCA and SCC, with >90% MHC-I HC loss in 12% of AdenoCA and 22% of SCC tumors (**figure 1A,D**). At the other extreme, 16 AdenoCAs (17%) and 10 SCCs (17%) had MHC-I expression in 100% of cells. We observed heterogeneity in MHC-I HC expression within each tumor, as CT and PT cores of the same tumor often displayed distinct MHC-I HC expression patterns (**figure 1A,D**). HC10^32^ reacts with unfolded HLA-A, -B and -C heavy chains, so it may not identify HLA Class I molecules folded with β2m; thus, we analyzed the correlation of β2m with HLA I HC expression. HLA HC and β2m staining were significantly correlated in AdenoCA and SCC (**figure 1B,E**), suggesting that low HC10 staining may be related to decreased surface expression of folded MHC-I dimers as well. This finding was consistent at the gene expression level (**supplemental figure S3**). Nonetheless, variability in MHC-I HC for each β2m classification suggested other regulators of MHC-I expression and/or antigen presentation may have affected MHC-I loss in Cohort 1. MHC-I HC loss thus is heterogeneously manifest across NSCLC patient tumors, and these may develop or progress differently in defined TME regions within individual patients.

**Figure 1.**
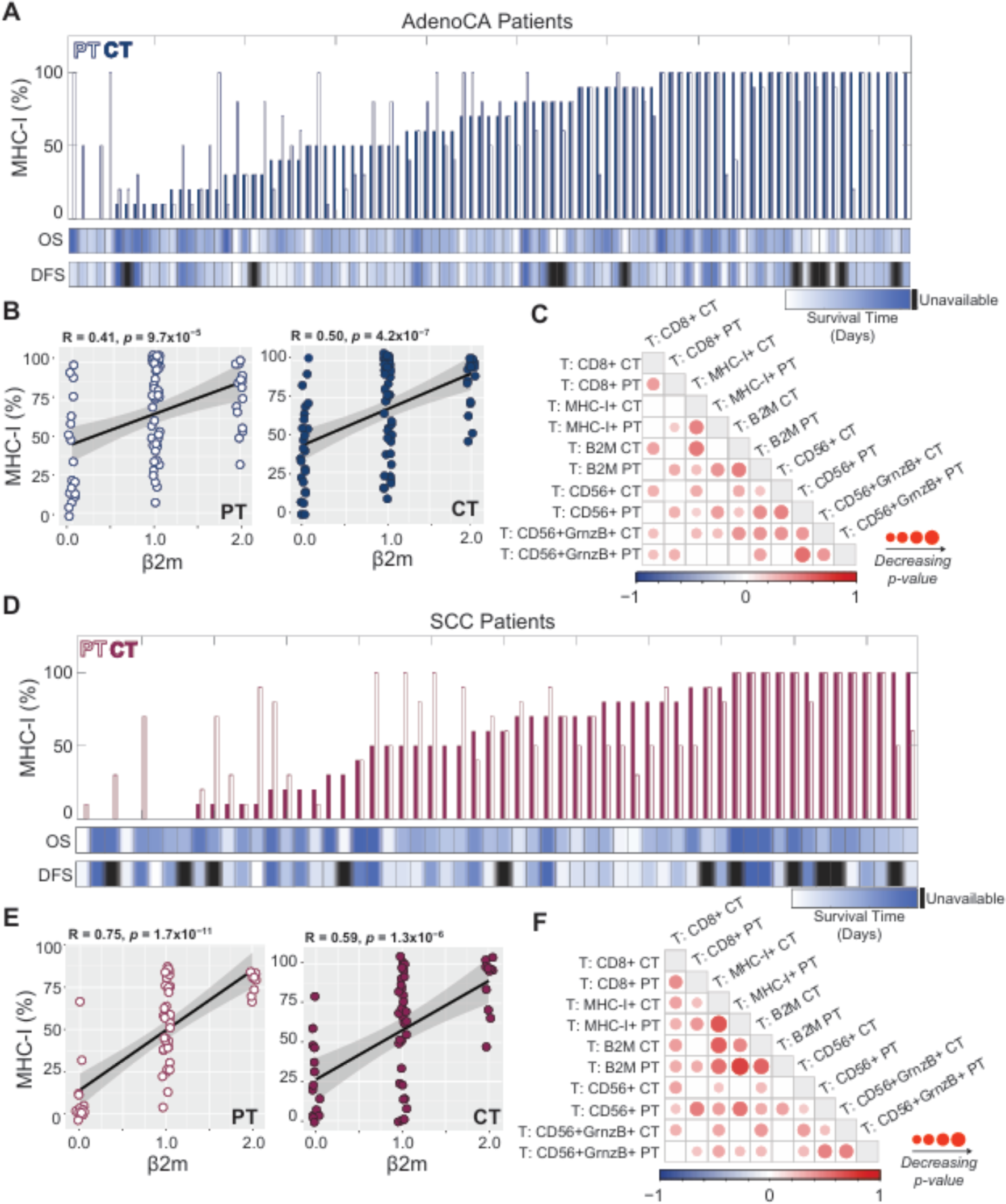
MHC-I expression exhibits intraand inter-tumor heterogeneity and correlates with lymphocyte occupancy. **(A&D)** MHC-I is variably expressed in tumors and across AdenoCA (A) and SCC (**D**) patients. (**B&E**) β2M expression correlates positively with MHC-I expression in AdenoCA (**B**) and SCC (**E**). (**C&F**) The heatmaps show Spearman correlations (R) among IHC features in AdneoCA (**C**) and SCC (**F**). Insignificant correlations (*p*>0.05) are shaded white; identity correlations are shaded grey.

Given its significant loss in NSCLC, we asked whether tumor cell MHC-I expression may be associated with NK and CD8 T cell infiltrates into the TME. In pairwise Spearman correlations of IHC features across all patient tumors, we found that the density of CD56^+^ and CD8^+^ cells significantly correlated with expression levels of both MHC-I HC and β2m in AdenoCA and SCC (**figure 1C,F**). Moreover, CD56^+^ cells correlated with CD8^+^ cells, most profoundly in PT regions.

### Concurrent tumor-infiltrating CD8^+^ and CD56^+^ immune cells are associated with patient survival in human NSCLC independent of MHC-I expression

Prior work showed that the density of tumor infiltrating CD8 T cells is associated with better patient survival.^15^ ^16^ Given MHC-I’s effect on target cell recognition by CD8 T cells and NK cells, we asked if MHC-I HC expression was associated with patient outcome. We did not detect any significant association between DFS or OS and MHC-I HC in CT, PT, or averaged across CT and PT (**figure 2A-B, supplemental figure S4**). We thus investigated whether densities of intra-tumoral CD8^+^ or CD56^+^ cells were associated with DFS or OS outcomes for Cohort 1 patients (**figure 2C-F**). High intra-tumoral CD56^+^ and CD8^+^ cell counts averaged across CT and PT were each positively associated with DFS and OS (**figure 2C-F**). Similar trends were observed when considering CT and PT separately (**supplemental figures S5-S6**). Further assessment revealed that simultaneously high CD8^+^ and CD56^+^ counts were more strongly associated with prolonged OS (**figure 2H**). Their joint impact on DFS was less striking, but evident when considering density of CD8^+^ and CD56^+^ cells in CT only (**figure 2G, supplemental figure S7**). Observations from Kaplan-Meier analyses were consistent when adjusting for patient age and sex in a Cox proportional hazards model (HR=0.199, *p*<1×10^-3^), and statistical power was insufficient to adjust for cancer stage and prior chemotherapy treatment (**supplemental table 1**). Collectively, these data demonstrate that combined presence of NK cells and CD8 T cells predicts better OS, pointing towards potential complementary antitumor activities.

**Figure 2.**
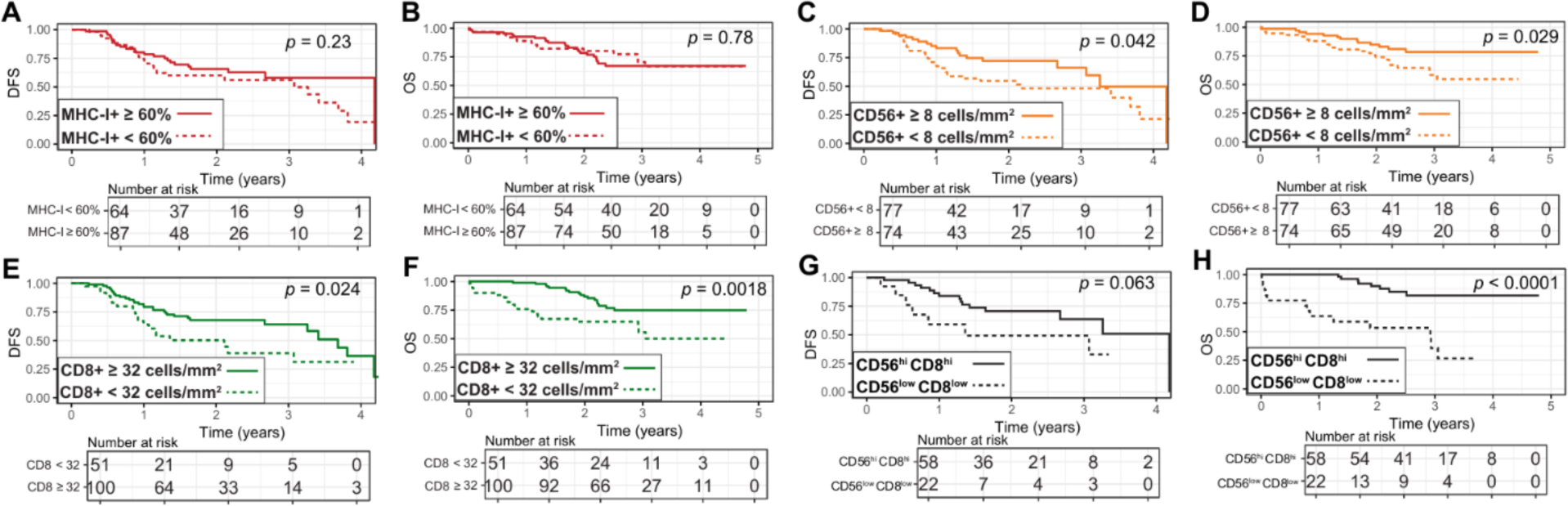
Survival probability increases with increasing CD8^+^ and CD56^+^ lymphocyte infiltrates. Kaplan-Meier analysis predicts DFS or OS with respect to tumor cell MHC-I HC expression **(A&B)**, CD56^+^ cell counts **(C&D)**, CD8^+^ cell counts **(E&F)**, or coincident CD56^+^ and CD8^+^ counts **(G&H)** averaged between CT/PT. *P*-value, log-rank test.

### mIF reveals the extent of MHC-I expression variation in patient resected tumor tissues

To further explore the relationship between lymphocyte infiltration and activation and tumor cell MHC-I expression considering its vast heterogeneity in NSCLC and the increased prognostic power observed with simultaneous infiltration of NK cells and CD8 T cells, we implemented mIF imaging with combined spatial analysis of whole tumor sections.

Like the TMA analysis, we observed variable MHC-I HC expression by mIF (**figure 3A**). Because prior work demonstrated that CD8 T cell infiltrations in NSCLC are heterogeneous across tumor regions,^19^ we developed a tumor regional approach to delineate spatial relationships between NK cells, T cells and tumor cell MHC-I HC expression (**supplemental figure S1, Methods**). Cohort-wide MHC-I expression per region revealed total MHC-I HC loss in 26% of regions and >90% loss in 47% of regions, with similar distributions observed in AdenoCA and SCC (**figure 3B**). Wide-ranging inter- and intra-patient MHC-I HC expression variability was observed with the mean expression per patient ranging from 0.1%-81% of panCK^+^ cells, and at least one region per tumor displaying >30% loss (**figure 3C**). Tumor cell MHC-I HC loss was thus common among NSCLC patients and heterogeneous across tumor resections.

**Figure 3.**
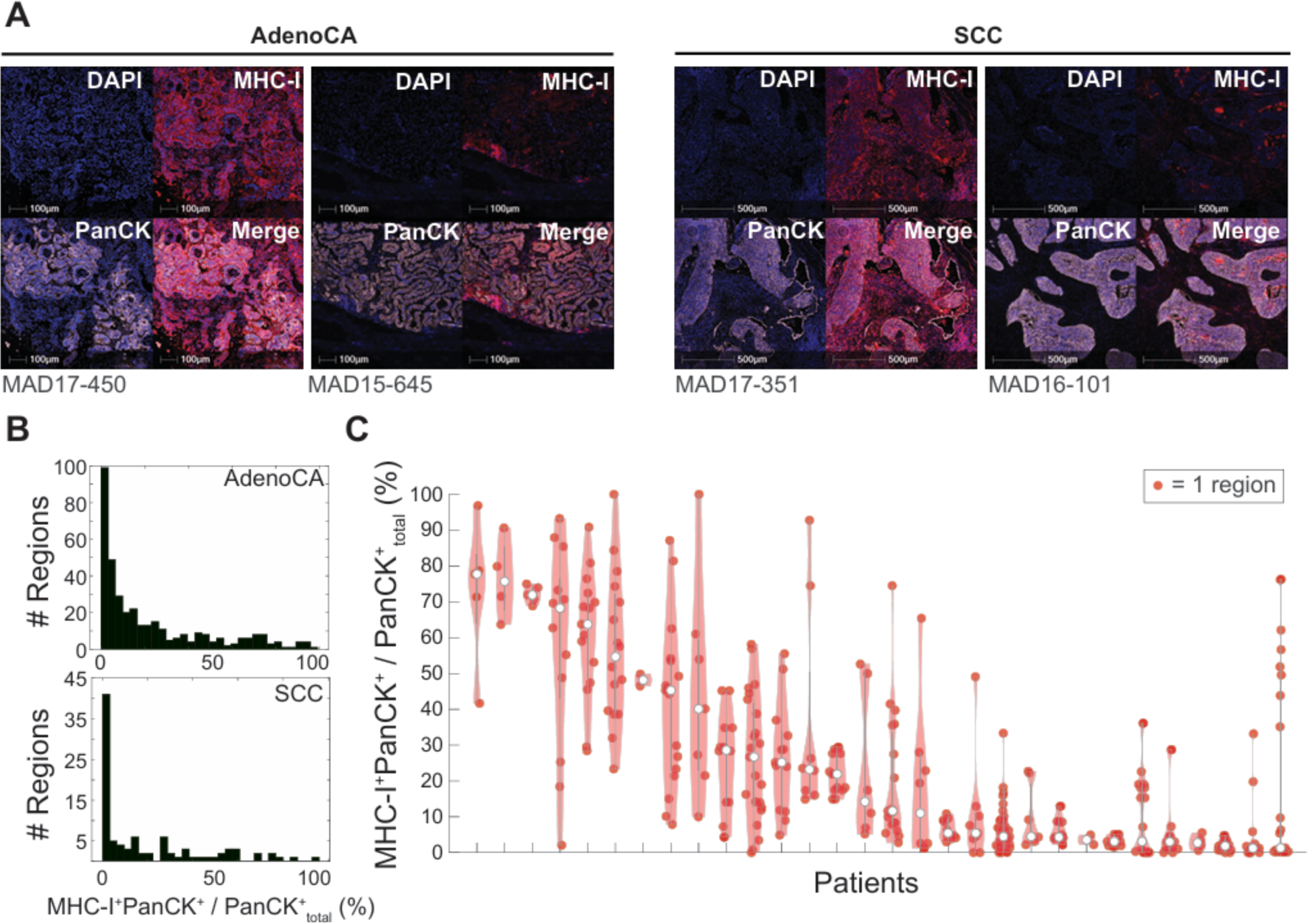
mIF imaging reveals extent of MHC-I HC expression variation in resected patient tumor tissues. **(A)** Representative images exemplify MHC-I loss in AdenoCA and SCC. **(B)** Histograms of (%) MHC-I^+^ tumor cells in each region of AdenoCA and SCC. **(C)** The (%) MHC-I^+^ tumor cells in each region per patient for AdenoCA and SCC.

### Tumor cell MHC-I deficiency is associated with decreased NK and T cell presence and activity in the NSCLC TME

We next investigated how NK cell, or CD8 T cell infiltration and activation corresponded to tumor cell MHC-I HC variability by building OPLSDA models to compare the presence of T cells and NK cells in MHC-I-disparate AdenoCA (**figure 4A-D**) or SCC (**figure 4E-H**) tumor regions. Lymphocyte activation was inferred by IFNγ staining in all Cohort 2 resections, though IFNγ is a secreted cytokine and as such its source cannot be definitively determined by mIF. Three tumors displayed strong IFNγ staining in nests lacking substantial lymphocyte infiltration, possibly indicative of a non-lymphocyte source of IFNγ. Nonetheless, in the remaining tumors (n=33) we observed IFNγ predominantly in the presence of NK cells and T cells, indicative of lymphocyte activation (**figure 5, supplemental figure S8**).

**Figure 4.**
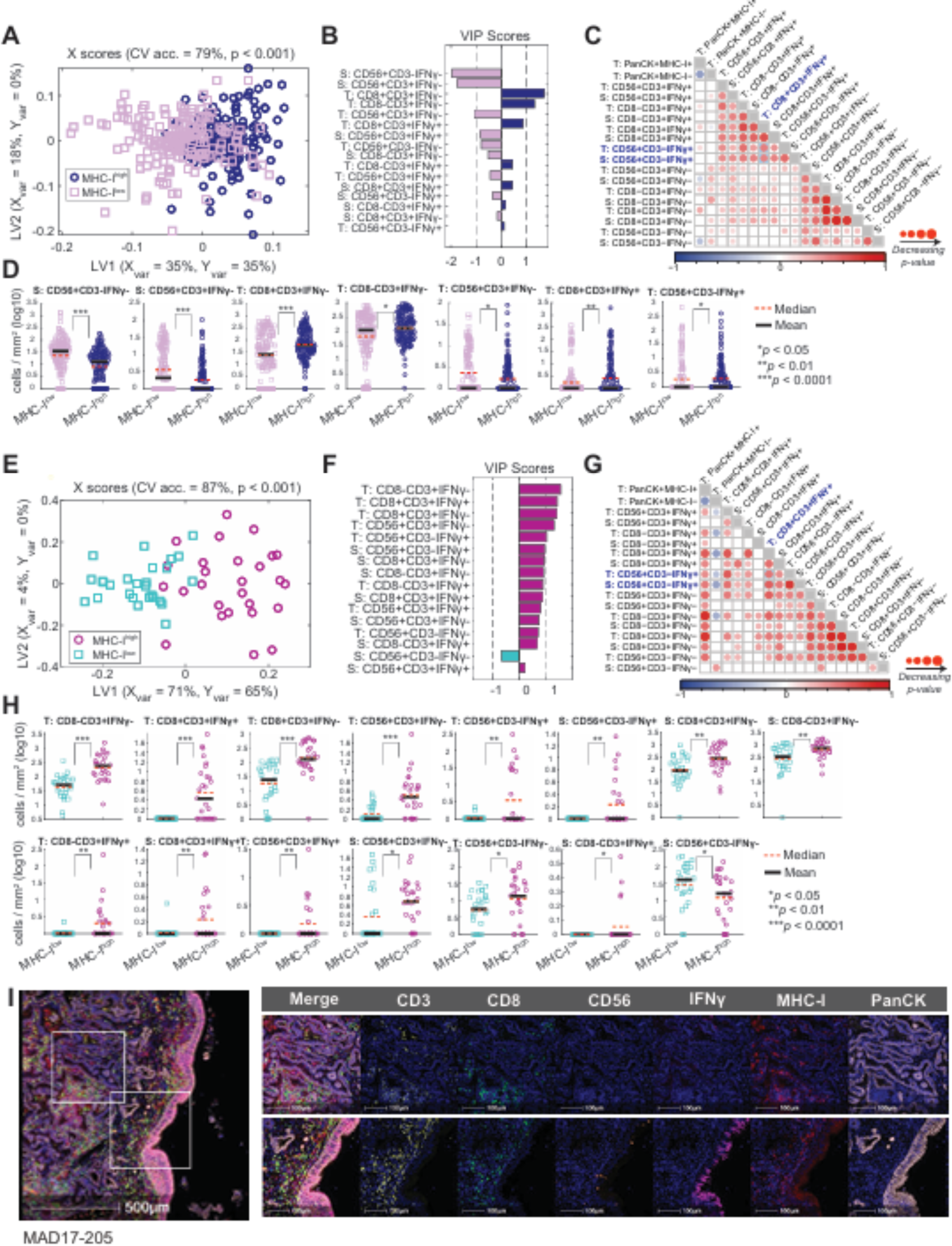
Tumor infiltrated lymphocytes and IFNγ expression are associated with high tumor cell MHC-I expression. OPLSDA models discriminate between tumor regions with MHC-I^+^ tumor cell densities (cells/mm^2^) above or below the median in AdenoCA **(A-D)** and SCC **(E-H)**. CV, cross-validation accuracy. Significance was determined by a permutation test (*p*<0.001). **(A&E)** X scores plot, where each point represents one region projected onto latent variables 1 and 2 (LV1&LV2). **(B&F)** VIP scores are shown artificially oriented in the direction of loadings on LV1. |VIP|>1 indicates variables with greater than average influence on the separation between groups. **(C&G)** Spearman correlations among immunologic features in tumor (T) and stroma (S). Insignificant correlations (*p*>0.05) are shaded white, identity correlations are shaded grey. **(D&H)** Univariate comparisons between model features. Wilcoxon rank sum test with Benjamini-Hochberg correction (\**p*<0.05; \*\**p*<0.01; \*\*\**p*<0.0001). Only features with *p*_adj_<0.05 are shown. **(I)** Representative image shows variation in lymphocyte occupancy and IFNγ staining in MHC-I^−^ (top) or MHC-I^+^ (bottom) regions of the same tumor.

**Figure 5.**
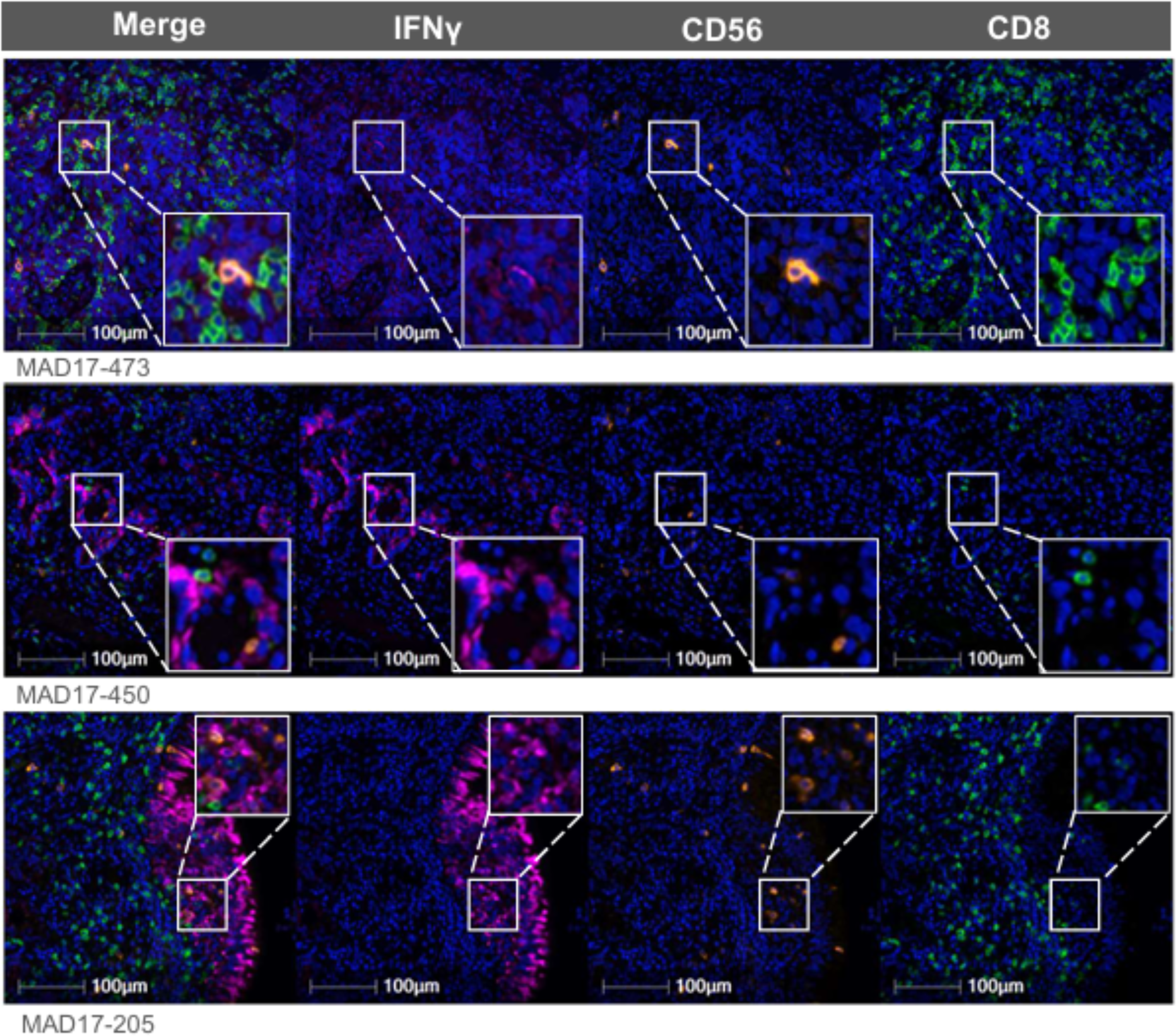
NK cells and CD8 T cells colocalize in areas with IFNγ staining. Representative images show NK cells and CD8 T cells clustered near IFNγ in 3 patient tumors.

The OPLSDA models were able to discriminate between tumor regions with disparate MHC-I expression based on lymphocyte infiltrates with 79% (AdenoCA) or 87% (SCC) accuracy (**figure 4A,E**). VIP scores from the AdenoCA model revealed that stromal CD3^−^CD56^+^ NK cells as well as both stroma- and tumor-associated CD56^+^CD3^+^ cells were enriched in tumor regions with tumor MHC-I expression below the median, whereas tumor-associated CD3^+^CD8^−^ (putative CD4^+^) and CD3^+^CD8^+^ T cells were enriched in regions with MHC-I expression above the median (|VIP|>1) (**figure 4B**). Tumor regions with high MHC-I expression also displayed a greater density of IFNγ^+^ CD8 T cells and NK cells (**figure 4D**), and Spearman correlation analysis revealed that tumoral IFNγ^+^ CD8 T cell densities were positively correlated with IFNγ^+^ NK cells in the stroma and tumor nest (**figure 4C,G**).

VIP scores from the SCC model indicated that CD3^+^CD8^−^ and CD3^+^CD8^+^ T cells, CD3^−^ CD56^+^ NK cells, and CD56^+^CD3^+^ cells were among the top features contributing to the separation between MHC-I replete and deficient tumors (**figure 4F,H**). Akin with the AdenoCAs, the only lymphocyte population found enriched in MHC-I low SCC tumor regions were IFNγ^−^ stromal NK cells. We observed a clear lack of immune infiltration into tumor nests with MHC-I loss in SCC tumors (**supplemental figure S9**). Additionally, we observed significant negative pairwise correlations between MHC-I^−^ tumor cell counts and IFNγ^+^ NK cells, CD56^+^CD3^+^ cells, CD3^+^CD8^+^ T cells, and CD3^+^CD8^−^ T cells, whereas IFNγ^−^ stromal NK cells showed a positive correlation further distinguishing their presence in SCC with MHC-I downregulation (**figure 4G**). Together, these data indicate that IFNγ-expressing NK cells and T cells coinhabited and were associated with tumor regions expressing high MHC-I (**figure 4I**).

### Local neighborhood analysis of NK cells and CD8 T cells in the TME reveals tightly clustered lymphocytes marked by IFNγ staining near MHC-I^+/–^ tumor cells

Given the observed differences in lymphocyte presence in MHC-I-disparate NSCLC tumors, we questioned whether IFNγ^+^ NK cells and CD8 T cells differ from IFNγ^−^ lymphocytes with respect to their proximity to other immune cells and MHC-I expressing tumor cells. To uncover phenotypic differences in cellcell communication between activated NK cells and CD8 T cells, we determined the cellular neighborhood profile of these effector cell types using spatial, cell-centric analyses of the mIF images (**supplemental figure S2**). The cellular neighborhood profile scores of both tumor- and stroma-associated cells were input into an OPLSDA model to delineate colocalization differences between IFNγ^+/–^ NK cells (**figure 6A-C**) and IFNγ^+/–^ CD8 T cells (**figure 6E-G**). Both models accurately distinguished between IFNγ^+/–^ lymphocytes based on their neighborhood profiles with cross-validation accuracy scores of 96% and 97%, respectively (**figure 6A,E**). Comparing the two models, we observed that both IFNγ^+^ NK cells and CD8 T cells associated with all IFNγ^+^ lymphocyte subsets more than IFNγ^−^ cells (**figure 6B,F**). In agreement with our regional analyses (**figure 4**), we found that NK cells and CD8 T cells were more likely to be IFNγ^+^ in the tumor as opposed to the surrounding stroma (**figure 6D,H**). Moreover, we observed that IFNγ^+^ NK cells and IFNγ^+^ CD8 T cells in the tumor nests colocalized with MHC-I^+^ tumor cells more frequently than did IFNγ^−^ lymphocytes, suggesting preferential localization of activated cells to tumors with MHC-I expression (**figure 6D,H**). Strikingly, IFNγ^+^ NK cells and CD8 T cells were more frequently associated with MHC-I^−^ tumor cells compared to their IFNγ^−^ counterparts, indicating infiltration of IFNγ-secreting lymphocytes into both MHC-I^+^ and MHC-I^−^ tumor nests (**figure 6I**). To validate these observations over larger radii, we quantified the number of neighbors <200 μm from a center cell.^35^ We observed IFNγ^+^ CD8 T cells had fewer MHC-I^−^ neighbors than MHC-I^+^ neighbors, and their association with MHC-I^−^ tumor cells was less than expected by random chance (**figure 6J**). This may signify deliberate enrichment of IFNγ^+^ CD8 T cells in MHC-I-bearing tumors. In contrast, IFNγ^+^ NK cells associated similarly with MHC-I^+^ and MHC-I^−^ tumor cells, suggesting that NK cells may become or remain activated in the MHC-I-deficient TME. Marked colocalization of IFNγ^+^ NK cells with other IFNγ^+^ lymphocytes including NK cells, CD8^−^ T cells, and CD8^+^ T cells additionally suggests that lymphocytes may be activated *en masse* in MHC-I-bearing tumors. IFNγ produced in these clusters of activated lymphocytes thus may be indicative of type I immune responses in the TME.

**Figure 6.**
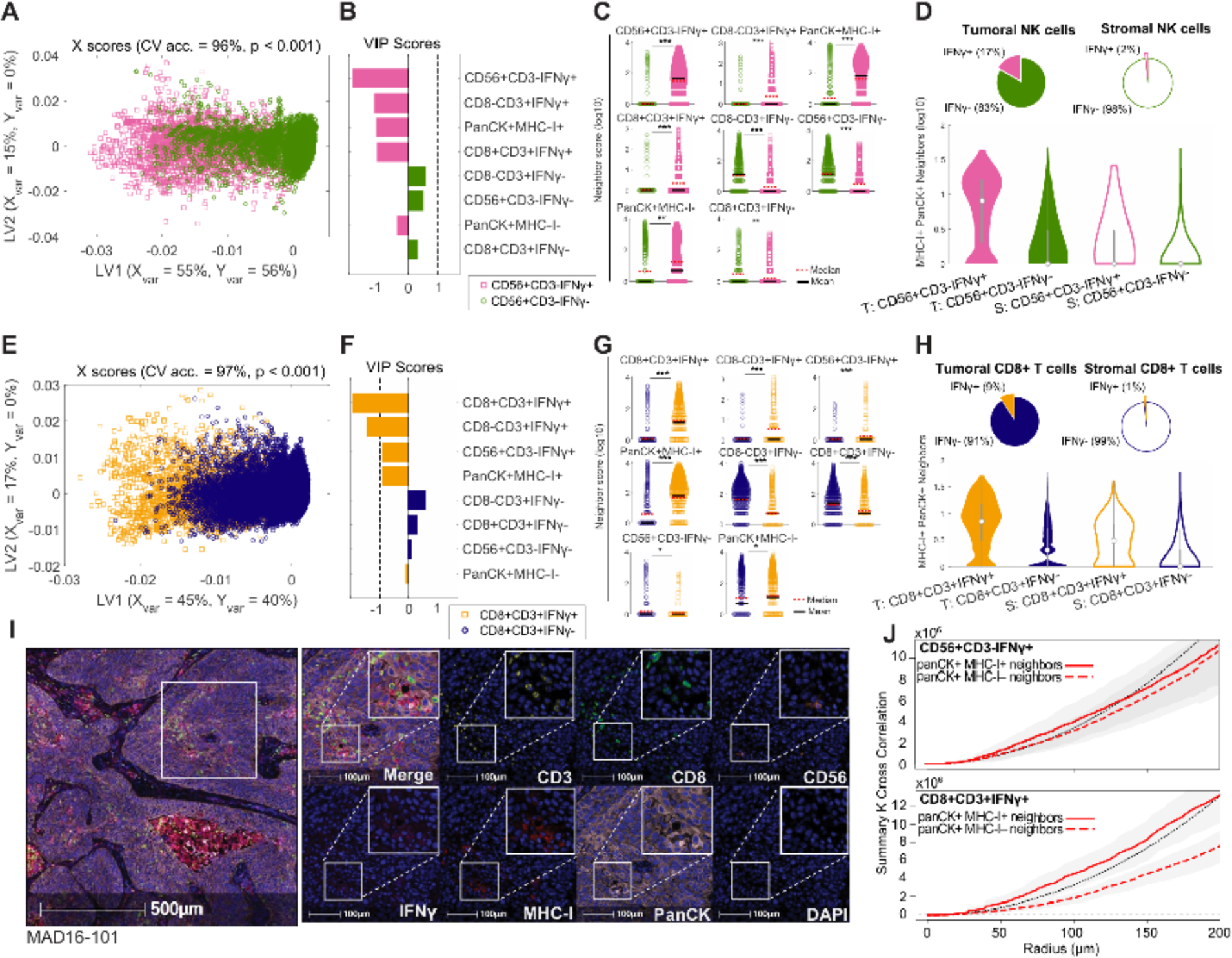
IFNγ^+^ NK cells and IFNγ^+^ CD8 T cells associate with other activated lymphocytes and MHC-I^+^ tumor cells. OPLSDA models discriminate IFNγ^+/–^ NK cells **(A-C)** and IFNγ^+/–^ CD8 T cells **(E-G)**. CV, cross-validation accuracy. Significance was determined by a permutation test (*p*<0.001). **(A&E)** X scores plot, where each point represents one region projected onto latent variables 1 and 2 (LV1&LV2). **(B&F)** VIP scores are shown artificially oriented in the direction of loadings on LV1. |VIP|>1 indicates a variable with greater than average influence on the separation between groups. **(C&G)** Univariate comparisons between model features. Wilcoxon rank sum test with Benjamini-Hochberg correction (\**p*<1×10^-27^; \*\**p*<1×10^-30^; \*\*\**p*<1×10^-50^). **(D&H)** The average counts of MHC-I^+^ tumor cell neighbors for IFNγ^+/–^ NK cells **(D)** or IFNγ^+/–^ CD8 T cells **(H)** in tumor (T) or stroma (S) regions. **(I)** Representative mIF image showing a cluster of lymphocytes colocalized with IFNγ in an MHC-I^−^ tumor nest. **(J)** The K function plotted against increasing radii for IFNγ^+^ NK cells and IFNγ^+^ CD8 T cells as the target cell and either MHC-I^+^ (red solid line) or MHC I^−^ (red dashed line) center cells. Grey shading, 95% confidence intervals. Black line, Poisson (null) distribution.

## DISCUSSION

While targeting immune checkpoints with CPI has had a substantive impact on the treatment of NSCLC, a minority of patients display durable responses. Whether this is due to hindered immune infiltration, functional exhaustion, tumor cell evasion of immune recognition by loss of MHC-I expression, or alternative mechanisms remains to be determined. Understanding the basis of this selectivity will increase the appropriate targeting of CPI or combination therapies to patients most likely to benefit and will ultimately lead to innovative approaches to expand the proportion of patients who receive clinical benefit.

We hypothesized that coordinated activities of both CD8 T cells and NK cells may be needed to mediate lysis of tumor cells while also preventing tumor cell escape via an MHC-I loss mechanism. We analyzed resected patient tumor tissues by both traditional IHC and mIF to simultaneously probe NK cells and CD8 T cells *in situ*, their effector activity indicated by IFNγ staining, and their co-localization in MHC-I HC disparate tumors. To define differences in lymphocyte activation and intercellular communication in MHC-I^+/–^ tumors, we leveraged the inherent heterogeneity of the TME by developing a suite of spatial analysis methods spanning whole tumors, tumor sub-regions, and single-cell enumeration of NK cell and T cell neighborhoods. This use of integrative systems consistently pointed towards coordinated anti-tumor activity of NK cells and T cells, particularly in MHC-I^+^ tumors.

Consistent with prior work,^8^ ^11^ ^13^ both cohorts displayed stark inter- and intra-tumoral MHC-I expression heterogeneity. Although MHC-I loss can lead to the development of CD8 T cell refractory tumors, we did not observe an association between tumor cell MHC-I HC expression and patient survival in this study. Others have speculated cohort-specific composition, clinicopathologic characteristics, genetic contribution and/or survival/events may obfuscate any potential association.^11–13^ MHC-I loss thus may indirectly impact patient survival by regulating lymphocyte occupancy and activation.

IHC analysis of Cohort 1 revealed that tumors with simultaneously high counts of CD8^+^ and CD56^+^ cells were more strongly associated with OS compared to tumors with high CD56^+^ or CD8^+^ cells only. Positive correlations between CD8^+^ and CD56^+^ cell counts suggests possible intercellular communication playing a role in tumor control. Using mIF to concurrently enumerate NK cell and CD8 T cell infiltrates in Cohort 2, we observed positively correlated tumor-infiltrating NK cells and CD8 T cells with greater densities in tumor nests expressing high levels of MHC-I, indicating coordinated infiltration. Furthermore, spatial analysis of local cellular neighborhoods revealed infiltrated NK and CD8 T cells are present in clusters marked by IFNγ activity together with CD8^−^ T cells. Though much less frequent, similar immune clusters were observed in tumors with MHC-I HC loss, suggesting that an effector response involving NK cells, CD8 T cells, and CD4 T cells can be active, and possibly leveraged, even in a setting considered refractory to lymphocyte immune control.

Whether intra-tumoral IFNγ promotes tumor cell MHC-I expression, immune recruitment and colocalization, or coordinated lymphocyte effector activity awaits further analysis, but these findings suggest that CPI plus novel approaches designed to jointly elicit NK and T cell effector activities may be key to developing next generation immunotherapies. Antigen-stimulated NK cells have been shown to provide support to CD8 T cell antitumor immunity in a mouse model of lung adenocarcinoma.^25^ Likewise, human induced pluripotent stem cell (iPSC-)derived NK cells have been shown to enhance T cell recruitment and activation to tumors, coinciding with improved T cell sensitivity to CPI and enhanced antitumor control.^38^ Cytokine activation can additionally invigorate NK cells to curb or regress tumor outgrowth, even in MHC-I^−^ tumors.^28^ The potential benefit of NK cells in NSCLC is bolstered by prior work that showed that human NK cell cytolytic activity is associated with lower cancer risk,^39^ whereas NK cell deficiency is associated with increased cancer risk.^40–42^ Our data contrast recent observations in soft tissue sarcoma that NK cell infiltrates indicate a poor prognosis and do not colocalize with CD8 T cells or MHC-I^+^ tumor cells, indicating that the role of NK cells may be context-specific.^43^

Our findings that NK cells infiltrated into tumor nests and displayed evidence of IFNγ activity in NSCLC begs the question, why might tumoral NK cells become ineffective drivers of anti-tumor immunity? Emerging evidence suggests that NK cell activity may be hindered in the lung cancer TME. Human NKp46^+^ NK cells were shown to infiltrate the invasive margin of lung tissue stroma surrounding tumor nests in NSCLC patients with no apparent effect on clinical outcomes ^24^. These intra-tumoral NK cells displayed inferior effector activities in comparison to their counterpart NK cells in peripheral blood.^24^ Whether due to suppressive mechanisms associated with the lung environment itself^44^ or more direct suppression via tumor cells remains unresolved as cancer cells can downregulate or secrete NK receptor ligands to evade natural killing, and the TME is rife with suppressive factors which can impede NK cell activity.^45^ Prior work in mouse models has further shown NK cells also undergo disarming in MHC-I^−^ tumors rendering them hyporesponsive and unable to mediate tumor cell clearance.^28^ ^46^ As our study affirms the significant association with survival and notable infiltration into MHC-I^+^ tumor nests of NK cells in NSCLC, future studies are needed to uncover the root causes of NK cell disarming in the lung cancer TME to fully realize their potential as anti-tumor effectors.

Though these multifaceted spatial analyses consistently revealed NK cell and CD8 T cell co-occurrence within NSCLC tumors, there are several limitations to consider. The TMA data from Cohort 1 were manually generated by a pathologist, a concern which is mitigated as automated analysis yielded similar results. Cohort 2 was limited by a relatively small sample size, warranting future investigations to verify the associations observed in this study. mIF imaging was limited to six phenotypic markers plus nuclear staining, thus constraining analysis of the full complexity of the tumor-associated immune milieu. Consequently, we relied on IFNγ expression as a marker of lymphocyte activity and were unable to test for expression of other cytotoxic markers and inhibitory ligands which may obscure true lymphocyte activity levels. As IFNγ is a secreted cytokine, we were unable to definitively determine its source; however, analysis of our data without three select tumors which displayed strong IFNγ staining in the absence of lymphocytes yielded similar results. The majority of IFNγ staining in the remainder of the cohort was colocalized with a heavy lymphocyte presence. Ongoing work with higher dimensional imaging (e.g., spatial transcriptomics) will provide a more comprehensive view of immune cell diversity, occupancy, proliferation, and activation in the TME. Because the cohort was composed of only complete resections, we were unable to assess tissues after chemotherapy or immunotherapy treatment. It remains to be determined whether NK cells drive recruitment and/or activation of CD8 T cells in the TME, or vice versa. Likewise, whether differences in lymphocyte populations and activity in tumors with MHC-I loss results from differences in lymphocyte infiltration, expansion, or retention in the TME remains unclear. Other outstanding questions include whether patient tumors are infiltrated by circulating NK cells with higher baseline cytotoxic activity, by lung resident NK cells with lower baseline cytotoxic activity, or both. Whether these different types of NK cells may affect other tumor-infiltrating immune cells, such as helper T cells or myeloid cells, and whether there are specific interactions that either cause or arise from the spatial associations observed in the data remains to be determined. Future mechanistic studies in animal models will help to delineate the effect of MHC-I on the balance of NK cells and their impact on the tumor-immune landscape.

Notwithstanding these challenges, we have shown that tumor cell MHC-I HC loss negatively impacted lymphocyte occupancy and activity in the NSCLC TME and that NK cell and CD8 T cells jointly infiltrate tumor nests which positively impacts patient survival. This evidence of lymphocyte activation and intercellular communication in the presence of MHC-I^+^ tumors highlights both the vital importance and promise for developing novel immunotherapies that leverage both NK cells and CD8 T cells to enhance immunologic control of NSCLC.

## Declarations

### Ethics approval and consent to participate

Not applicable.

### Consent for publication

Not applicable.

### Availability of data and material

Demographics data for Cohort 1 and 2 are included in Supplementary Spreadsheets 1 and 2, respectively. Algorithms developed for this study (O-PLSDA and cell neighborhood scoring) are available on GitHub (https://github.com/Dolatshahi-Lab). Quantitative IHC and mIF data are available as Supplementary Spreadsheets 3-4. IHC and mIF images will be made available by the authors upon reasonable request.

### Authors contributions

REW, SD and MGB wrote the manuscript with input from all authors. REW, NA, JMO, IM, KG, CL, RDG, CLS, TNJB, SD and MGB contributed to the conception, design, and development of the project. REW, NA, JMO, IM, KG, GFH, MH, and NW performed experiments and/or analyzed data. All authors approved the final version of the manuscript.

## Acknowledgements

This work was presented at the 2023 American Association of Immunologists (AAI) Annual Meeting (Abstract 62.08). We thank Dr. Patcharin Pramoonjago and outstanding technical support from the Biorepository and Tissue Research Facility (BTRF) which is supported by the University of Virginia (UVA) School of Medicine, Research Resource Identifiers (RRID): SCR_022971. This work was also supported by the UVA Molecular Immunologic & Translational Sciences (MITS) core facility and the co-pay by the UVA Cancer Center Support Grant number 2P30CA044579-26.

## List of abbreviations

CT: Central Tumor
CPI: Checkpoint Inhibitor Immunotherapy
HR: Hazards Ratio
HC: Heavy Chain
IFNγ: Interferon gamma
MHC-I: MHC class I
mIF: Multiplexed Immunofluorescence
NK cell: Natural Killer cell
NSCLC: Non-Small Cell Lung Cancer
SCC: Squamous Cell Carcinoma
OPLSDA: Orthogonalized Partial Least Squares Discriminant Analysis
PT: Peripheral Tumor
VIP: Variable Importance in Projection

## REFERENCES

1. Molina JR, Yang P, Cassivi SD, et al. Non-small cell lung cancer: epidemiology, risk factors, treatment, and survivorship. Mayo Clinic proceedings 2008;83(5):584–94. doi: 10.4065/83.5.584

2. Siegel RL, Miller KD, Fuchs HE, et al. Cancer statistics, 2022. CA: a cancer journal for clinicians 2022;72(1):7-33. doi: 10.3322/caac.21708 [published Online First: 2022/01/13]

3. Chen Z, Fillmore CM, Hammerman PS, et al. Non-small-cell lung cancers: a heterogeneous set of diseases. Nat Rev Cancer 2014;14(8):535–46. doi: 10.1038/nrc3775 [published Online First: 2014/07/25]

4. Travis WD, Brambilla E, Noguchi M, et al. International Association for the Study of Lung Cancer/American Thoracic Society/European Respiratory Society: international multidisciplinary classification of lung adenocarcinoma: executive summary. Proc Am Thorac Soc 2011;8(5):381–5. doi: 10.1513/pats.201107-042ST [published Online First: 2011/09/20]

5. Stankovic B, Bjorhovde HAK, Skarshaug R, et al. Immune Cell Composition in Human Non-small Cell Lung Cancer. Frontiers in immunology 2018;9:3101. doi: 10.3389/fimmu.2018.03101 [published Online First: 2019/02/19]

6. Garrido F, Aptsiauri N, Doorduijn EM, et al. The urgent need to recover MHC class I in cancers for effective immunotherapy. Current opinion in immunology 2016;39:44–51. doi: 10.1016/j.coi.2015.12.007 [published Online First: 2016/01/23]

7. Thunnissen E, van der Oord K, den Bakker M. Prognostic and predictive biomarkers in lung cancer. A review. Virchows Arch 2014;464(3):347–58. doi: 10.1007/s00428-014-1535-4 [published Online First: 2014/01/15]

8. Redondo M, Ruiz-Cabello F, Concha A, et al. Altered HLA class I expression in non-small cell lung cancer is independent of c-myc activation. Cancer research 1991;51(9):2463–8.

9. Korkolopoulou P, Kaklamanis L, Pezzella F, et al. Loss of antigen-presenting molecules (MHC class I and TAP-1) in lung cancer. British journal of cancer 1996;73(2):148–53. doi: 10.1038/bjc.1996.28 [published Online First: 1996/01/01]

10. Delp K, Momburg F, Hilmes C, et al. Functional deficiencies of components of the MHC class I antigen pathway in human tumors of epithelial origin. Bone Marrow Transplant 2000;25 Suppl 2:S88–95. doi: 10.1038/sj.bmt.1702363 [published Online First: 2000/08/10]

11. Ramnath N, Tan D, Li Q, et al. Is downregulation of MHC class I antigen expression in human non-small cell lung cancer associated with prolonged survival? Cancer immunology, immunotherapy : CII 2006;55(8):891–9. doi: 10.1007/s00262-005-0085-7 [published Online First: 2005/09/28]

12. Li X, Guo F, Liu Y, et al. NLRC5 expression in tumors and its role as a negative prognostic indicator in stage III non-small-cell lung cancer patients. Oncol Lett 2015;10(3):1533–40. doi: 10.3892/ol.2015.3471 [published Online First: 2015/12/02]

13. Datar IJ, Hauc SC, Desai S, et al. Spatial Analysis and Clinical Significance of HLA Class-I and Class-II Subunit Expression in Non-Small Cell Lung Cancer. Clinical cancer research : an official journal of the American Association for Cancer Research 2021;27(10):2837–47. doi: 10.1158/1078-0432.CCR-20-3655 [published Online First: 2021/02/20]

14. Kikuchi E, Yamazaki K, Torigoe T, et al. HLA class I antigen expression is associated with a favorable prognosis in early stage non-small cell lung cancer. Cancer Sci 2007;98(9):1424–30. doi: 10.1111/j.1349-7006.2007.00558.x [published Online First: 2007/07/25]

15. Geng Y, Shao Y, He W, et al. Prognostic Role of Tumor-Infiltrating Lymphocytes in Lung Cancer: a Meta-Analysis. Cellular physiology and biochemistry : international journal of experimental cellular physiology, biochemistry, and pharmacology 2015;37(4):1560–71. doi: 10.1159/000438523 [published Online First: 2015/10/30]

16. Zeng DQ, Yu YF, Ou QY, et al. Prognostic and predictive value of tumor-infiltrating lymphocytes for clinical therapeutic research in patients with non-small cell lung cancer. Oncotarget 2016;7(12):13765–81. doi: 10.18632/oncotarget.7282 [published Online First: 2016/02/13]

17. Schalper KA, Brown J, Carvajal-Hausdorf D, et al. Objective measurement and clinical significance of TILs in non-small cell lung cancer. J Natl Cancer Inst 2015;107(3) doi: 10.1093/jnci/dju435 [published Online First: 2015/02/05]

18. Esendagli G, Bruderek K, Goldmann T, et al. Malignant and non-malignant lung tissue areas are differentially populated by natural killer cells and regulatory T cells in non-small cell lung cancer. Lung cancer 2008;59(1):32–40. doi: 10.1016/j.lungcan.2007.07.022

19. Obeid JM, Wages NA, Hu Y, et al. Heterogeneity of CD8(+) tumor-infiltrating lymphocytes in non-small-cell lung cancer: impact on patient prognostic assessments and comparison of quantification by different sampling strategies. Cancer immunology, immunotherapy : CII 2017;66(1):33–43. doi: 10.1007/s00262-016-1908-4 [published Online First: 2016/10/23]

20. Federico L, McGrail DJ, Bentebibel SE, et al. Distinct tumor-infiltrating lymphocyte landscapes are associated with clinical outcomes in localized non-small-cell lung cancer. Annals of oncology : official journal of the European Society for Medical Oncology 2022;33(1):42–56. doi: 10.1016/j.annonc.2021.09.021 [published Online First: 2021/10/16]

21. Guillerey C, Huntington ND, Smyth MJ. Targeting natural killer cells in cancer immunotherapy. Nat Immunol 2016;17(9):1025–36. doi: 10.1038/ni.3518

22. Malmberg KJ, Carlsten M, Bjorklund A, et al. Natural killer cell-mediated immunosurveillance of human cancer. Semin Immunol 2017;31:20–29. doi: 10.1016/j.smim.2017.08.002

23. Vivier E, Raulet DH, Moretta A, et al. Innate or adaptive immunity? The example of natural killer cells. Science 2011;331(6013):44–9. doi: 331/6013/44 [pii] 10.1126/science.1198687 [published Online First: 2011/01/08]

24. Platonova S, Cherfils-Vicini J, Damotte D, et al. Profound coordinated alterations of intratumoral NK cell phenotype and function in lung carcinoma. Cancer research 2011;71(16):5412–22. doi: 10.1158/0008-5472.CAN-10-4179

25. Schmidt L, Eskiocak B, Kohn R, et al. Enhanced adaptive immune responses in lung adenocarcinoma through natural killer cell stimulation. Proc Natl Acad Sci U S A 2019;116(35):17460–69. doi: 10.1073/pnas.1904253116 [published Online First: 2019/08/15]

26. Liu RB, Engels B, Arina A, et al. Densely granulated murine NK cells eradicate large solid tumors. Cancer research 2012;72(8):1964–74. doi: 10.1158/0008-5472.CAN-11-3208

27. Nicolai CJ, Wolf N, Chang IC, et al. NK cells mediate clearance of CD8(+) T cell-resistant tumors in response to STING agonists. Science immunology 2020;5(45) doi: 10.1126/sciimmunol.aaz2738 [published Online First: 2020/03/22]

28. Ardolino M, Azimi CS, Iannello A, et al. Cytokine therapy reverses NK cell anergy in MHC-deficient tumors. J Clin Invest 2014;124(11):4781–94. doi: 10.1172/JCI74337

29. Bottcher JP, Bonavita E, Chakravarty P, et al. NK Cells Stimulate Recruitment of cDC1 into the Tumor Microenvironment Promoting Cancer Immune Control. Cell 2018;172(5):1022–37 e14. doi: 10.1016/j.cell.2018.01.004

30. Kyrysyuk O, Wucherpfennig KW. Designing Cancer Immunotherapies That Engage T Cells and NK Cells. Annu Rev Immunol 2023;41:17–38. doi: 10.1146/annurev-immunol-101921-044122 [published Online First: 2022/11/30]

31. Erdag G, Schaefer JT, Smolkin ME, et al. Immunotype and immunohistologic characteristics of tumor-infiltrating immune cells are associated with clinical outcome in metastatic melanoma. Cancer research 2012;72(5):1070–80. doi: 10.1158/0008-5472.CAN-11-3218 [published Online First: 2012/01/24]

32. Stam NJ, Spits H, Ploegh HL. Monoclonal antibodies raised against denatured HLA-B locus heavy chains permit biochemical characterization of certain HLA-C locus products. J Immunol 1986;137(7):2299–306. [published Online First: 1986/10/01]

33. Mauldin IS, Mahmutovic A, Young SJ, et al. Multiplex Immunofluorescence Histology for Immune Cell Infiltrates in Melanoma-Associated Tertiary Lymphoid Structures. Methods Mol Biol 2021;2265:573–87. doi: 10.1007/978-1-0716-1205-7_40 [published Online First: 2021/03/12]

34. Contal C, O’Quigley J. An application of changepoint methods in studying the effect of age on survival in breast cancer. Computational Statistics & Data Analysis 1999;30(3):253–70. doi: 10.1016/S0167-9473(98)00096-6

35. Baddeley A, Turner R. spatstat: An R Package for Analyzing Spatial Point Patterns. Journal of Statistical Software 2005;12:1–42. doi: doi.org/10.18637/jss.v012.i06

36. Dolatshahi S, Butler AL, Siedner MJ, et al. Altered Maternal Antibody Profiles in Women With Human Immunodeficiency Virus Drive Changes in Transplacental Antibody Transfer. Clinical infectious diseases : an official publication of the Infectious Diseases Society of America 2022;75(8):1359–69. doi: 10.1093/cid/ciac156 [published Online First: 2022/03/05]

37. Grace PS, Dolatshahi S, Lu LL, et al. Antibody Subclass and Glycosylation Shift Following Effective TB Treatment. Frontiers in immunology 2021;12:679973. doi: 10.3389/fimmu.2021.679973 [published Online First: 2021/07/23]

38. Cichocki F, Bjordahl R, Gaidarova S, et al. iPSC-derived NK cells maintain high cytotoxicity and enhance in vivo tumor control in concert with T cells and anti-PD-1 therapy. Science translational medicine 2020;12(568) doi: 10.1126/scitranslmed.aaz5618 [published Online First: 2020/11/06]

39. Imai K, Matsuyama S, Miyake S, et al. Natural cytotoxic activity of peripheral-blood lymphocytes and cancer incidence: an 11-year follow-up study of a general population. Lancet 2000;356(9244):1795–9. doi: 10.1016/S0140-6736(00)03231-1

40. Orange JS. Natural killer cell deficiency. The Journal of allergy and clinical immunology 2013;132(3):515–25. doi: 10.1016/j.jaci.2013.07.020

41. Moon WY, Powis SJ. Does Natural Killer Cell Deficiency (NKD) Increase the Risk of Cancer? NKD May Increase the Risk of Some Virus Induced Cancer. Frontiers in immunology 2019;10:1703. doi: 10.3389/fimmu.2019.01703 [published Online First: 2019/08/06]

42. Mace EM, Orange JS. Emerging insights into human health and NK cell biology from the study of NK cell deficiencies. Immunol Rev 2019;287(1):202–25. doi: 10.1111/imr.12725 [published Online First: 2018/12/20]

43. Cruz SM, Sholevar CJ, Judge SJ, et al. Intratumoral NKp46(+) natural killer cells are spatially distanced from T and MHC-I(+) cells with prognostic implications in soft tissue sarcoma. Frontiers in immunology 2023;14:1230534. doi: 10.3389/fimmu.2023.1230534 [published Online First: 2023/08/07]

44. Marquardt N, Kekalainen E, Chen P, et al. Human lung natural killer cells are predominantly comprised of highly differentiated hypofunctional CD69(-)CD56(dim) cells. The Journal of allergy and clinical immunology 2017;139(4):1321–30 e4. doi: 10.1016/j.jaci.2016.07.043

45. Granzin M, Wagner J, Kohl U, et al. Shaping of Natural Killer Cell Antitumor Activity by Ex Vivo Cultivation. Frontiers in immunology 2017;8:458. doi: 10.3389/fimmu.2017.00458

46. Marcus A, Mao AJ, Lensink-Vasan M, et al. Tumor-Derived cGAMP Triggers a STING-Mediated Interferon Response in Non-tumor Cells to Activate the NK Cell Response. Immunity 2018;49(4):754–63 e4. doi: 10.1016/j.immuni.2018.09.016 [published Online First: 2018/10/18]

