## Supplemental Information for "Spatial colocalization and combined survival benefit of natural killer and CD8 T cells despite profound MHC class I loss in non-small cell lung cancer"

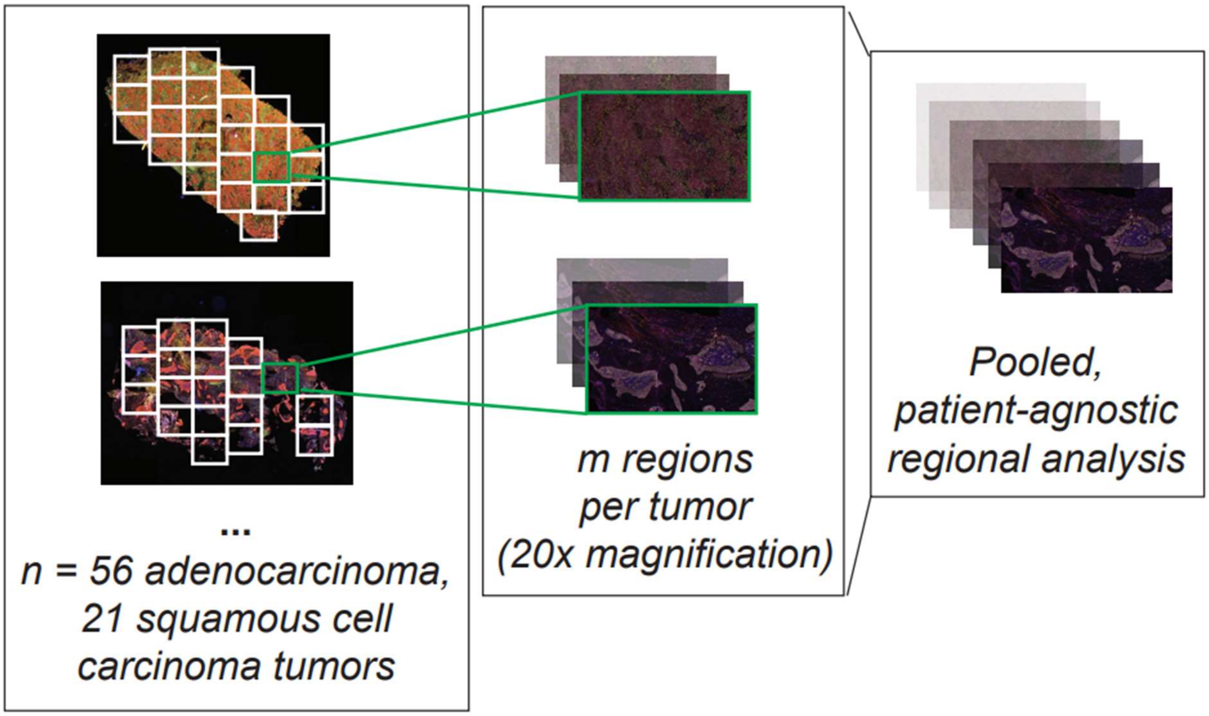


**Supplemental Figure S1. Regional analysis of resected tumors.** The schematic demonstrates systematic partitioning of tumor sections into sub-regions for more granular analysis. Left: Whole tumor slides are partitioned into 3 $mm^2$ regions. Center: each region is scanned at 20x magnification. Right: Regions are pooled across all tumors in a patient-agnostic analysis. In total, 394 regions across 29 adenocarcinoma tumors and 108 regions across 7 SCC tumors were collected. Regions with evident tumor cell CD56 expression or fewer than 50 cells enumerated were excluded. The remaining 361 adenocarcinoma regions and 89 SCC regions were analyzed. To validate that over-representation of tumors with a greater number of images did not bias our results, we iteratively sampled a random subset of images with equal representation from all tumors and observed similar results.

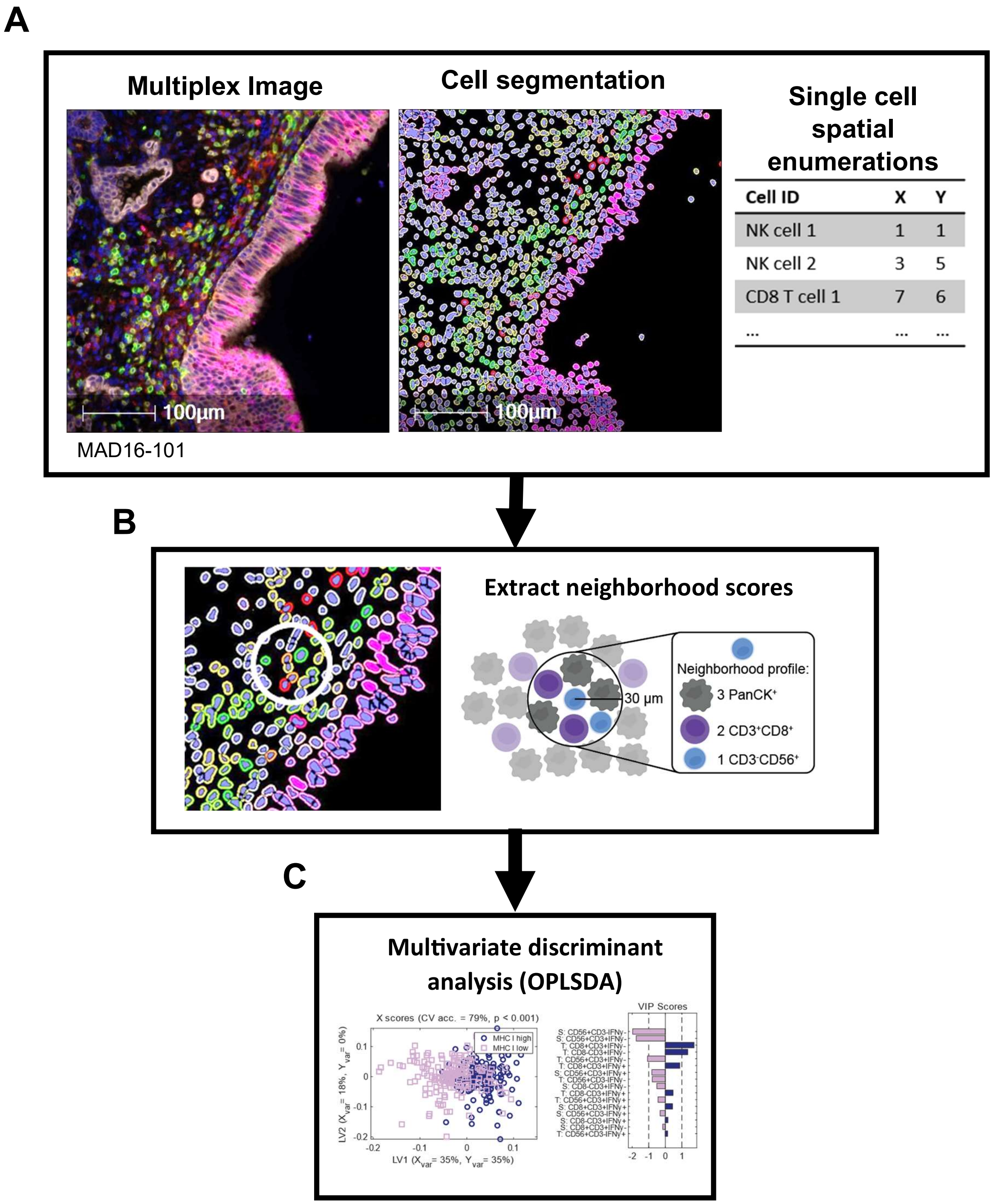


**Supplemental Figure S2.** Cell-centric determination of neighborhood profile. The diagram demonstrates the workflow of the neighborhood colocalization scoring algorithm developed in-house. Individual cells were segmented from tumors using Halo software. (A) Cells were classified by their expression of canonical markers and the spatial coordinates were saved for each cell. (B) These data were input into an algorithm which calculates the number of neighbor cells of each cell phenotype within a radius (<30 µm) of the center cell. (C) The neighborhood profiles were then input into multivariate discriminant analysis models to distinguish neighborhood profiles of two cell types of interest.

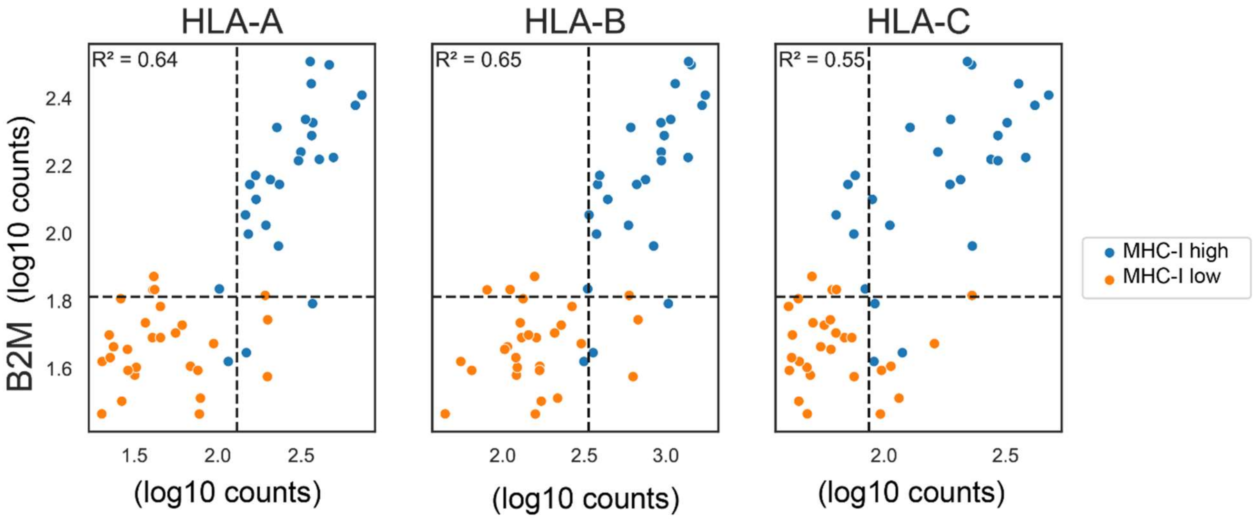


**Supplemental Figure S3: MHC class I subunit expression corresponds with HC10 staining classifications.** Counts (log10 transformed) of HLA-A, HLA-B, and HLA-C correlated with B2M in a subset of tumor cores from Cohort 1 (including both AdenoCA and SCC patients). Patients were colored by MHC class I high vs low groups as classified by HC10 staining.

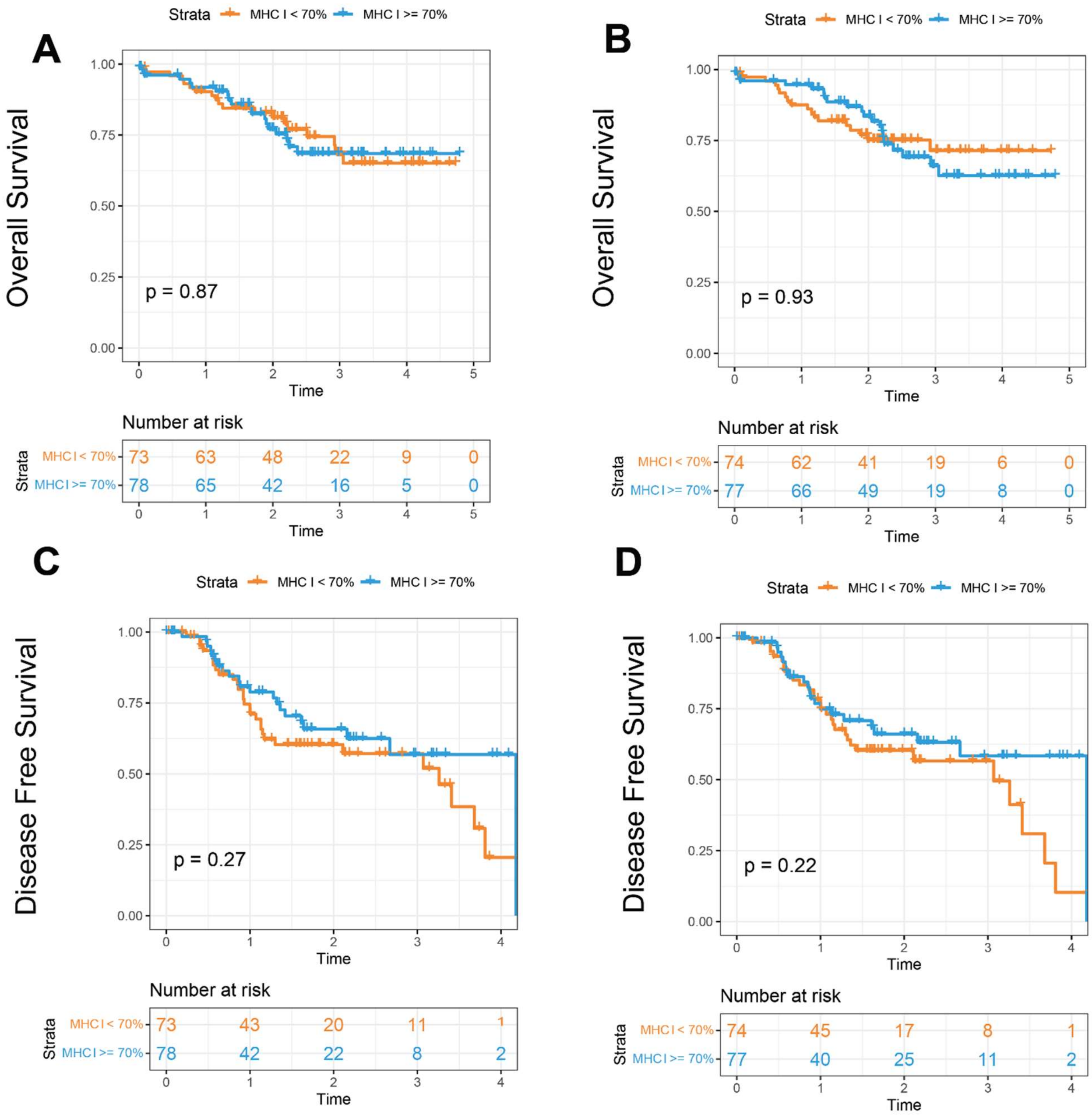


**Supplemental Figure S4. Survival probability by MHC-I HC expression.** The probability of OS (A-B) and DFS (C-D) in Cohort 1 was determined for patients with MHC-I expression cells above or below the cohort median. MHC-I was considered in central tumor (CT) (A,C) or peripheral tumor (PT) (B,D). Significance was determined by a log-rank test.

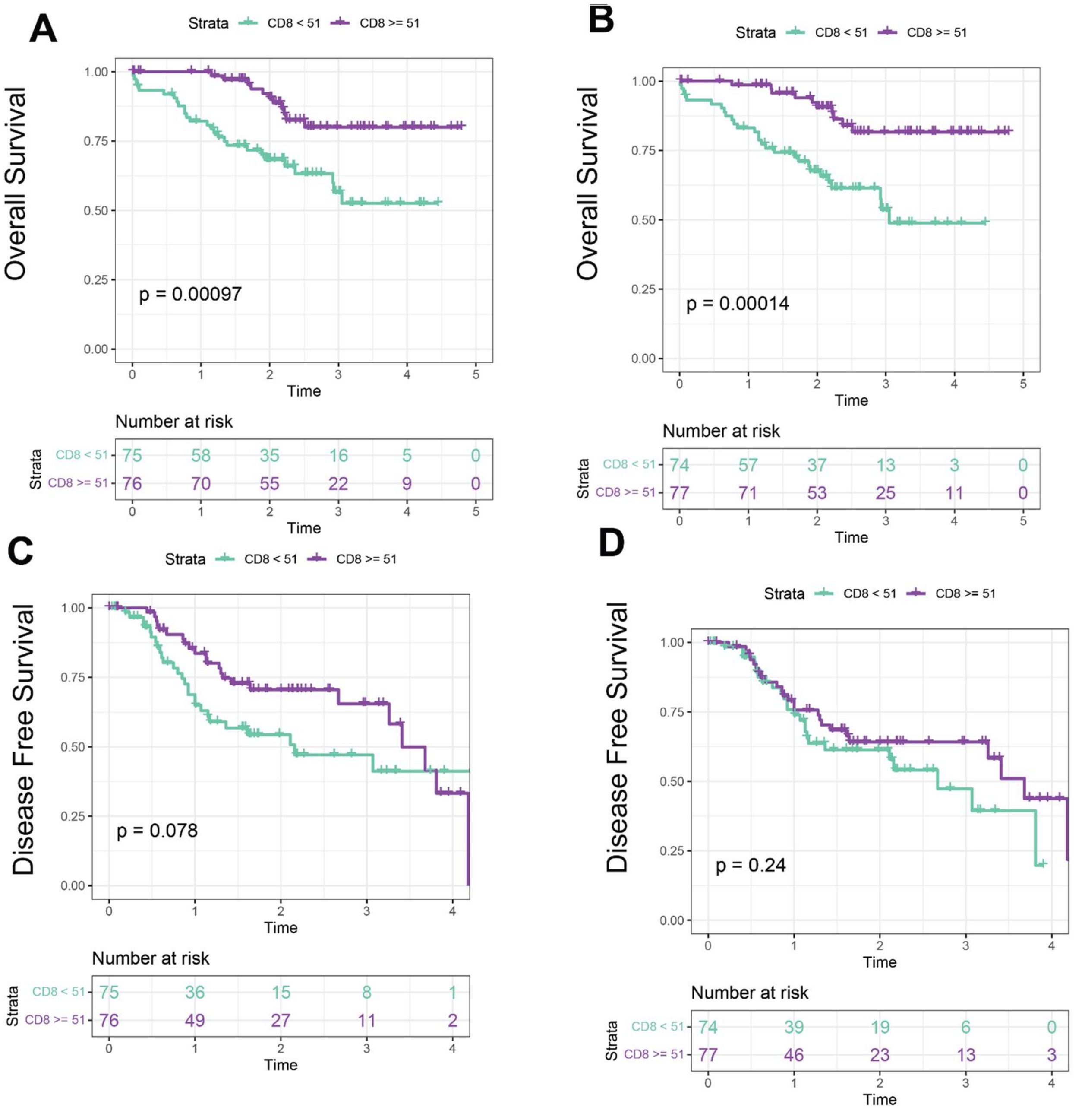


**Supplemental Figure S5. Survival probability by CD8+ counts.** The probability of OS (A-B) and DFS (C-D) in Cohort 1 was determined for patients with intra-tumor counts of $CD8^+$ cells above or below the cohort median. Cell counts were considered in central tumor (CT) (A,C) or peripheral tumor (PT) (B,D). Significance was determined by a log-rank test.

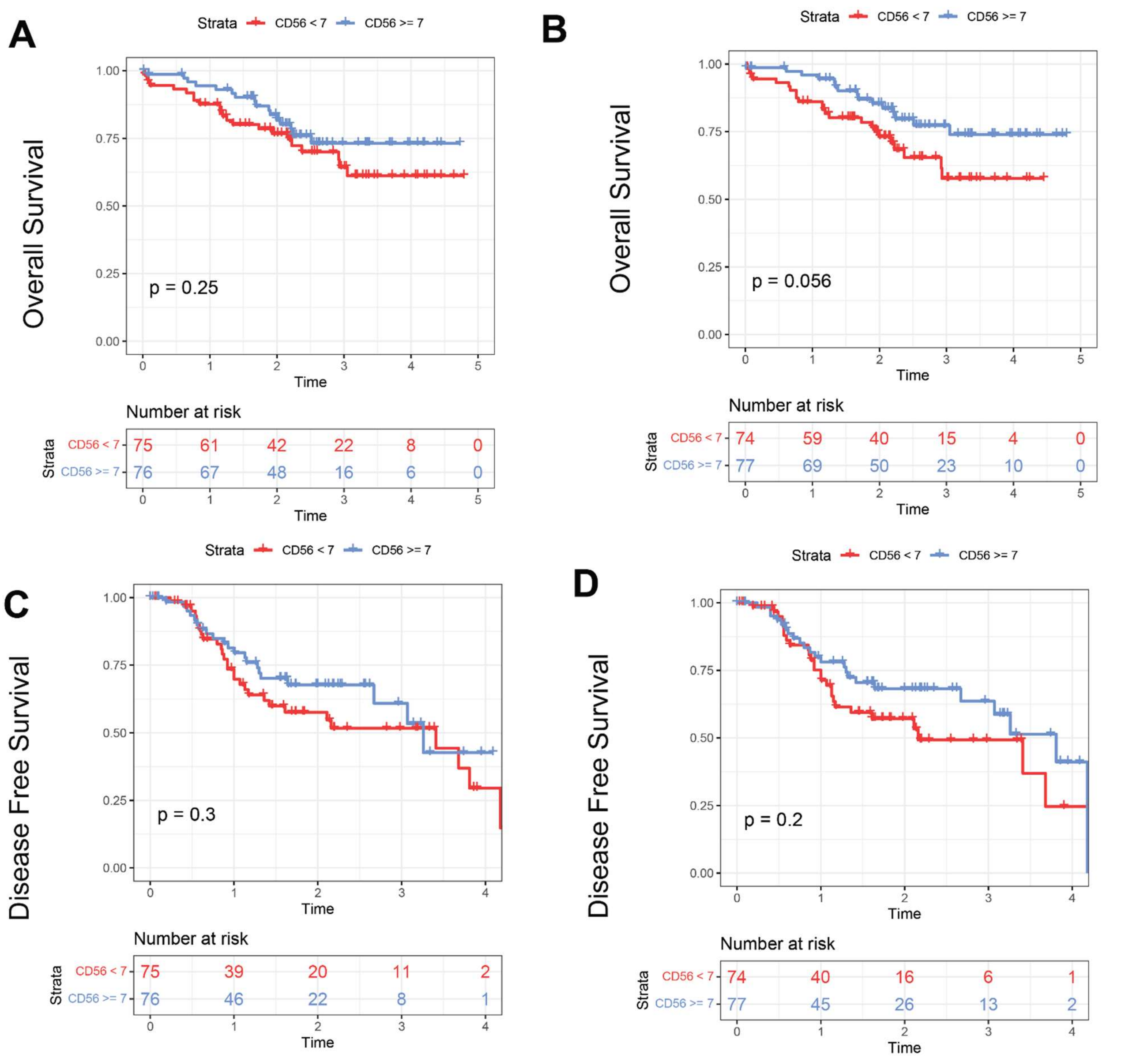


**Supplemental Figure S6. Survival probability by CD56+ counts.** The probability of OS (A-B) and DFS (C-D) in Cohort 1 was determined for patients with intra-tumor counts of $CD56^{+}$ cells above or below the cohort median. Cell counts were considered in central tumor (CT) (A,C) or peripheral tumor (PT) (B,D). Significance was determined by a log-rank test.

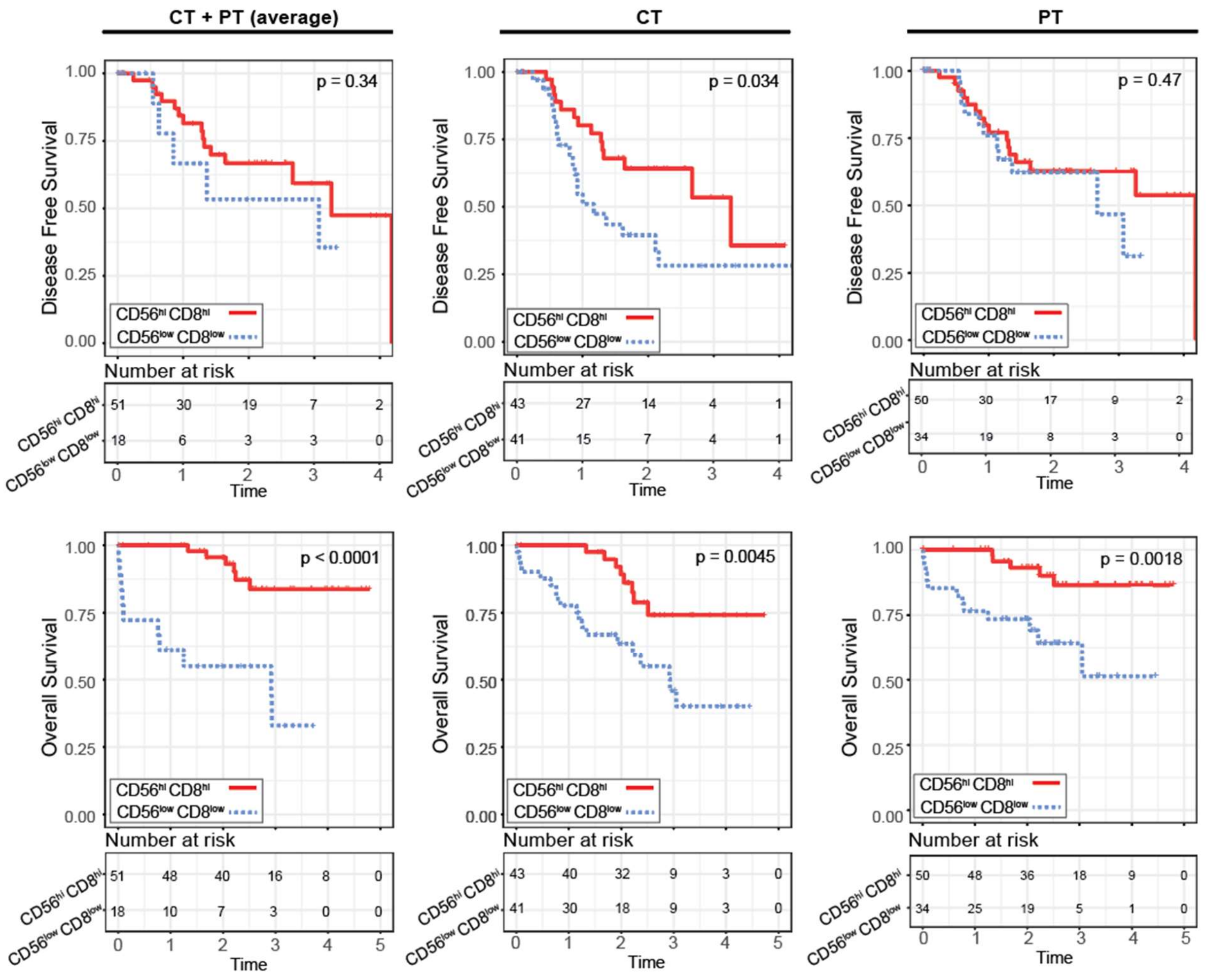


**Supplemental Figure S7. Coincident high NK and CD8 T cells increases survival probability.** The probability of disease free survival (top row) and overall survival (bottom row) in Cohort 1 was determined for patients with intra-tumor counts of both $CD8^+$ and $CD56^+$ cells above or below the cohort median cell count. Cell counts were either averaged across central and peripheral tumor (CT + PT average, left), considered for central tumor only (CT, middle), or peripheral tumor only (PT, right). Patients with missing values for one of these features were omitted from respective analyses. Significance was determined by a log-rank test.

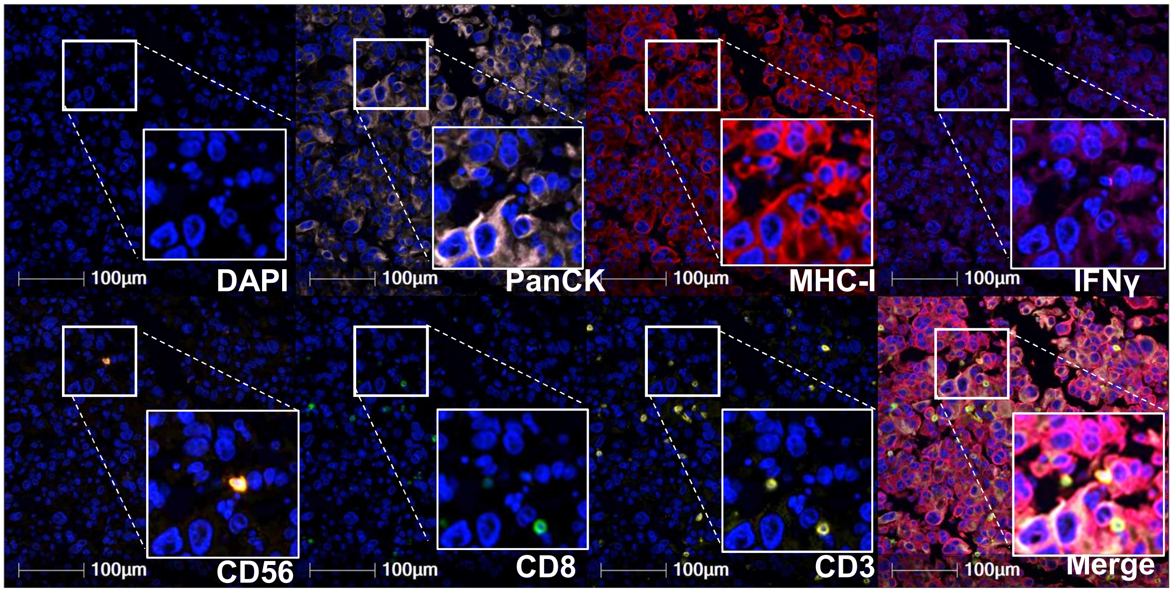


MAD16-180_[51355,16388]: ADENOCARCINOMA

**Supplemental Figure S8: Sample IFNγ+ NK cell in an MHC-I bearing tumor nest.**

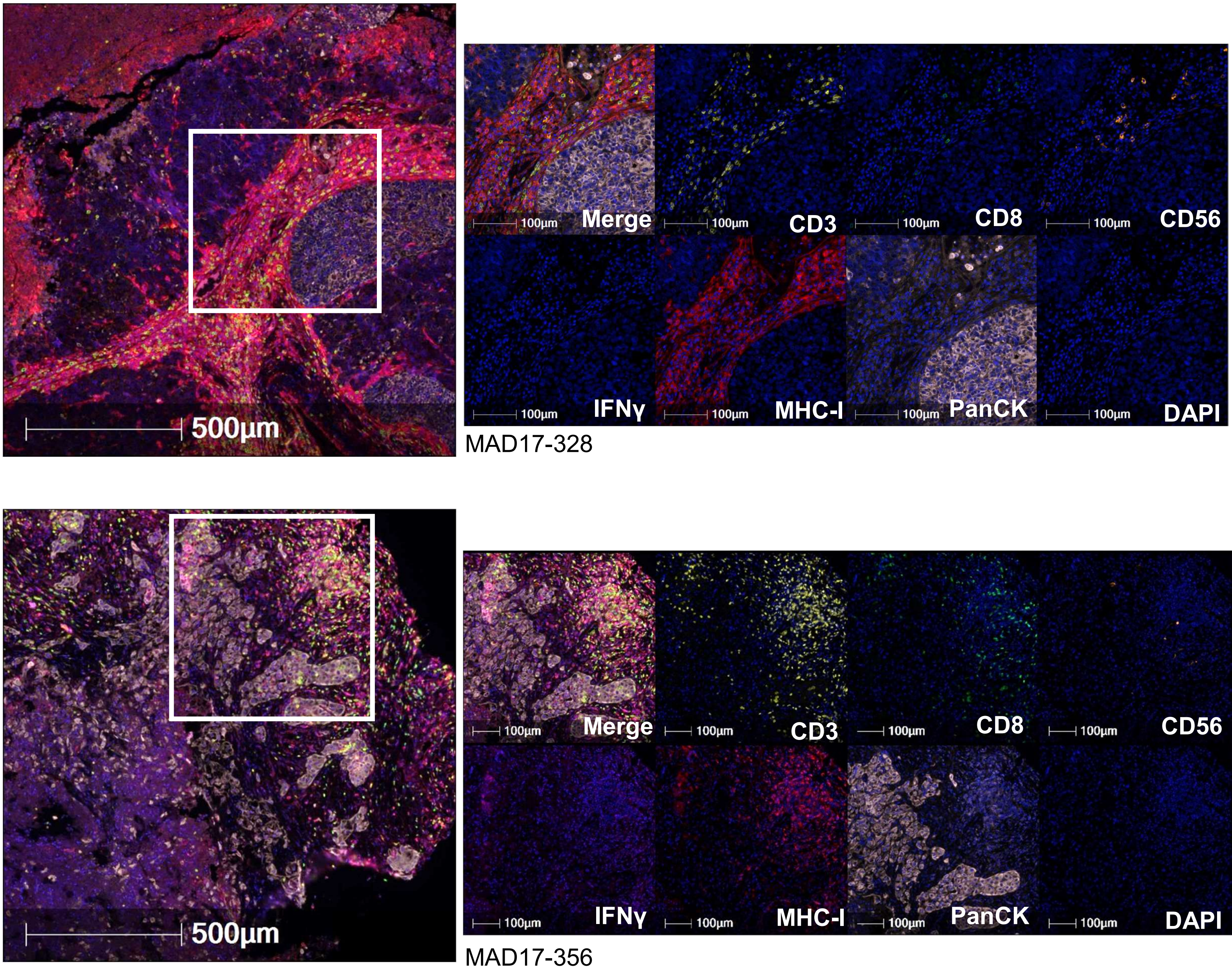


**Supplemental Figure S9: Stark absence of NK cell and T cell infiltration into MHC-I⁻ tumor nests.**

**Supplemental Table 1.** Multivariate Cox proportional hazards modeling was performed using R software version 4.2.1. Proportional hazards assumptions were tested using Schoenfeld residual analysis for univariate and multivariate analysis using a level of significance of 0.01. Multivariate Cox proportional hazards models assessed the relationship between survival endpoints (OS and DFS) and NK cell, CD8 T cell abundances, and MHC-I expression, after adjusting for age and sex. Prior treatment and stage were considered for inclusion in the models but were ultimately excluded because there were too few observations in some categories of these variables. Associations were considered significant for two-sided p values ≤ 0.05.

### MHCI+ Multivariate Cox proportional hazards model (OS)

| | HR [confint] | p |
|---|---|---|
| MHCI+ (>60%) | 0.97 [0.508, 1.87] | 0.9391 |
| Age | 1.04 [1.004, 1.08] | 0.0301 |
| Sex | 0.65 [0.333,1.27] | 0.2046 |

### MHCI+ Multivariate Cox proportional hazards model (DFS)

| | HR [confint] | p |
|---|---|---|
| MHCI+ (>60%) | 0.78 [0.44, 1.39] | 0.399 |
| Age | 0.98 [0.95,1.02] | 0.301 |
| Sex | 0.63 [0.35,1.13] | 0.12 |

### CD8 - Multivariate Cox proportional hazards model (OS)

| | HR [confint] | p |
|---|---|---|
| CD8 (>32) | 0.39 [0.206, 0.745] | **0.0043** |
| Age | 1.103 [0.998, 1.07] | 0.0664 |
| Sex | 0.64 [0.3293,1.24] | 0.1824 |

### CD8 - Multivariate Cox proportional hazards model (DFS)

| | HR [confint] | P |
|---|---|---|
| CD8 (>32) | 0.467 [0.26, 0.84] | **0.011** |
| Age | 0.97 [0.94, 1.01] | 0.1175 |
| Sex | 0.59 [0.33,1.05] | 0.0742 |

### CD56 - Multivariate Cox proportional hazards model (OS)

| | HR [confint] | p |
|---|---|---|
| CD56 (>8) | 0.496 [0.253, 0.971] | **0.0406** |
| Age | 1.04 [1.003, 1.08] | 0.0361 |
| Sex | 0.67 [0.3438,1.39] | 0.2341 |

### CD56 - Multivariate Cox proportional hazards model (DFS)

| | HR [confint] | p |
|---|---|---|
| CD56 (>8) | 0.58 [0.32, 1.03] | 0.0636 |
| Age | 0.98 [0.95,1.01] | 0.2437 |
| Sex | 0.64 [0.36,1.15] | 0.1375 |

### $CD56^{hi}CD8^{hi}$ - Multivariate Cox proportional hazards model (OS)

| | HR [confint] | p |
|---|---|---|
| $CD56^{hi}CD8^{hi}$ | 0.199 [0.08, 0.488] | **0.0004** |
| Age | 1.04 [0.992, 1.100] | 0.0958 |
| Sex | 0.1.49 [0.2641,1.71] | 0.404 |

### $CD56^{hi}CD8^{hi}$ - Multivariate Cox proportional hazards model (DFS)

| | HR [confint] | p |
|---|---|---|
| $CD56^{hi}CD8^{hi}$ | 0.437 [0.175, 1.09] | 0.0765 |
| Age | 0.965 [0.912,1.02] | 0.222 |
| Sex | 0.563 [0.225,1.407] | 0.2188 |
